# Three Dimensional Dynamics of Epithelial Monolayers

**DOI:** 10.64898/2026.03.10.710903

**Authors:** Silja Borring Låstad, Nigar Abbasova, Thomas Combriat, Dag Kristian Dysthe

## Abstract

Cells in epithelial monolayers are commonly quantified using a projected area and implicitly assuming constant cell volume and prism-shaped cells. These 2.5D assumptions ignore the volume and mass dynamics that may accompany density fluctuation in confluent space-filling tissues but remain inaccessible to the labelling- and intensity-based imaging techniques used to date. Here, we use time-lapse quantitative phase imaging (QPI) to obtain spatially and temporally resolved maps of height, volume, and dry mass in MDCK epithelial monolayers under physiological conditions. Three findings follow. (i) Cellular dry mass concentration is maintained to within ∼ 4.5 % across the monolayer and over time, even during large-amplitude pulsations, ruling out fluid (water) transport as the dominant driver of height and area dynamics. (ii) The mean height of the monolayer increases from ∼ 5.5 to ∼ 9 µm and the mean cell volume decreases by ∼ 25 % as the cell density doubles, evidence of contact inhibition of cell *size* rather than constant cell volume. Cell heights vary up to 30%, oscillate with a ∼ 5 h period, and follow gamma-shaped distributions. (iii) At cellular scales, both segmented-cell tracking and continuum mass-flux analysis show that projected cell volume is not conserved; mass conservation is recovered only after coarse-graining over ∼ 2 cell diameters and ∼ 1.6 h. Two non-exclusive mechanisms may explain this: non-prismatic cell geometry produces apparent volume fluctuations even at constant true volume, and periodic mass exchange between cells and the ECM produces genuine cyclic changes in cellular dry mass. A propagating elastic mode in the dry-mass structure factor (*c*_*s*_ ∼ 15 µm/h) further indicates that the monolayer behaves as a viscoelastic active medium rather than a plug-flow fluid. Each of these results requires simultaneous, time-resolved access to height and dry mass at the cellular scale and is therefore newly visible with QPI applied to epithelial monolayers. Together, they question the common 2.5D assumptions of prism-shaped cells, constant volume, plug-flow kinematics, and argue that quantitative continuum and cell-based models of epithelial dynamics must incorporate dry-mass-density regulation and 3D cell geometry.

## I. INTRODUCTION

Epithelial tissues are thin confluent monolayers that cover organs and line cavities, forming protective barriers with little extracellular matrix. Such tissues dynamically restructure to maintain organ integrity. Their collective cell motion underlies biological processes, including embryogenesis, morphogenesis, wound repair, and tumour metastasis [1–4].

Several modes of collective motion and shape change in epithelia are related to coordinated variations in cell size, but most studies quantify these dynamics with 2D measures of projected area and geometry. This ignores intrinsically 3D processes – height and volume changes – that accompany morphogenesis, epithelial–mesenchymal transitions, and density-dependent flows in confluent monolayers [5, 6]. Unlike active-matter systems with free space, epithelial sheets are space-filling, so density fluctuations necessarily entail reciprocal changes in cell height and/or volume. Time-lapse imaging reveals prominent area fluctuations [7], and recent evidence links these directly to volume dynamics implicating 3D cell size regulation in collective migration [8–10]. The observed area pulsations have been attributed to intercellular water flow [8, 9].

The use of 2D projections further presupposes a cell shape: that cells are prisms with vertical lateral membranes and equal apical and basal areas. However, non-prismatic shapes such as frustum and scutoids have been shown to be necessary in curved and invaginating epithelia [5, 11, 12], where they arise as a geometric consequence of tissue bending, yet flat confluent monolayers are still routinely modelled as prismatic. Recently, 3D models of epithelial layers have been proposed to give a more complete picture of epithelial dynamics [5, 13–16]. However, the scant experimental evidence is conflicting and a quantitative framework that connects volume fluctuations to tissue-scale motion remains missing, limiting our understanding of how mechanical forces and cell packing shape epithelial behaviour.

Changes in monolayer height are also related to the regulation of single-cell volume and mass. Size parameters (volume, dry mass, surface area) are homeostatically controlled and coupled across timescales [17, 18]. On fast scales (seconds to minutes), mechano-osmotic feedback and pump–leak regulation couple membrane/cortical tension to ion and water flux. On slower timescales (hours), metabolic regulation changes impermeable biomass, and cell volume adjusts accordingly to preserve intracellular concentrations (dry mass concentration), thereby coordinating growth with the cell cycle [17, 19–21]. However, the proposed volume-regulation pathways have not been systematically unified with the pulsatile, tissue-level behaviours characteristic of epithelial monolayers.

Here, we employ a label-free experimental technique that has not been used quantitatively for epithelial monolayers before. We report the use of time-lapse quantitative phase imaging (QPI) of Madin-Darby canine kidney (MDCK) monolayers, which yields spatially and temporally resolved maps of cell height, volume, and dry mass under physiological conditions. We make three main observations. First, the dry mass concentration is maintained within a narrow range across the monolayer and over time, even during large-amplitude pulsations of cell height and area. This constrains the role of intercellular water flow in driving the pulsations. Second, as the mean cell density slowly increases, the mean monolayer height increases, and the mean cell volume falls, indicating contact inhibition of cell size rather than constant cell volume. Third, we present what is, to our knowledge, the first direct test of plug-flow continuity equation that underlies 2.5D descriptions of monolayers, using direct measurements of the mass field: Both segmented-cell tracking and a continuum mass-flux analysis show that projected cell volume is not conserved at the scale of single cells. Mass conservation is recovered only after coarse-graining over about two cell diameters and 1.6 hours. This apparent non-conservation implies either that mass is exchanged locally and/or that the lateral cell membranes are not vertical.

## II. MATERIALS AND METHODS

We used label-free 2D QPI and 3D QPI (refractive index tomography) to study the evolution of cell height, cell volume, and cell dry mass under physiological conditions during MDCK monolayer pulsations. This relies on novel methods to determine the reference phase of 2D QPI data and the height of the cell layer in 3D QPI data. The complete description of data analysis and validation has recently been published separately [22].

### A. Measurement of mass, volume, and height

Cellular biomass, cell dry mass or cell nonaqueous mass, *m*_*d*_, is the total mass of everything in a cell except water and is dominated by macromolecules such as proteins (40–60%), nucleic acids (10–20%), lipids (5–20%), carbohydrates (5–20%), and small metabolites and ions (smaller, but non-negligible fraction) [23].

Few experimental techniques are available to measure the cell mass and volume of adherent cells in a petri dish. Confocal fluorescence microscopy has been used to estimate epithelial monolayer height [8–10, 18] with varying results, stimulated Raman microscopy can measure both protein and lipid mass [24], quantitative phase imaging (QPI) can measure total dry mass [25], refractive index tomography (3D QPI) can measure both total dry mass and cell volume [24, 26], and fluorescence exclusion measurement (fXm) can measure the volume of single cells [27].

The basis of QPI is the linear relation between refractive index *n* and dry mass concentration *c*_*d*_ = *m*_*d*_*/V*, where *m*_*d*_ is the dry mass in a volume *V* [28]:

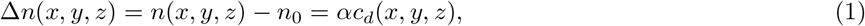

where *n*_0_ is the refractive index of the surrounding medium and the proportionality constant *α* is called the refraction increment. 3D QPI, or refractive index tomography, measures the 3D distribution of refractive indices of cells, *n*(*x, y, z*), and 2D QPI measures the integral of refractive index over the cells along the optical axis, ∫*n*(*x, y, z*) *dz*. Integrating over the volume of cell *i* and using an accepted value of the refractive increment, *α* = 0.18−0.2 ml/g [28–30] one obtains the general formula for dry mass of cell *i*:

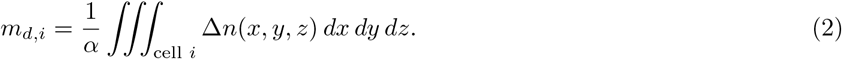

The dry mass, *m*_*d*_, measured by stimulated Raman microscopy, has been found to agree well with refractive index tomography measurements [24].

The height at every point in the image is determined differently in 2D and 3D QPI. In 3D QPI, the height *h*(*x, y*) is determined by a form of thresholding of the 3D refractive index *n*(*x, y, z*) in the *z*-axis [22]. In 2D QPI, the height is the integral of Δ*n* along the optical axis and inversely proportional to the mean dry mass concentration:

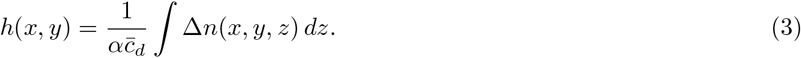

Segmentation of images is used to identify the area *A*_*i*_ occupied by each cell *i* in the monolayer. The 2.5D-approximation used for both 2D and 3D QPI assumes that the cell *i* fills the volume under the area *A*_*i*_ defined by the segmentation, and thus the cell volume is defined as *V*_*i*_ = *A*_*i*_*h*_*i*_, where *h*_*i*_ is the average height of the cell *i*. The general formula (2) for the dry mass of the cell *i* is, in the 2.5D-approximation evaluated as follows:

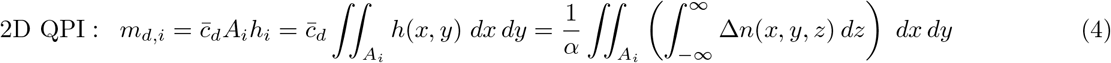

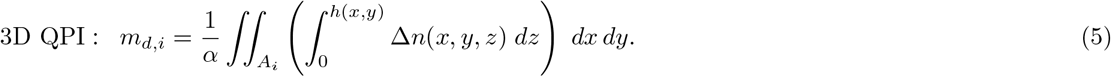

### B. QPI measurements and analysis

2D QPI was performed with Holomonitor M3 (Phase Holographic Imaging AB, Sweden). Height maps *h*(*x, y*) were exported from the Holomonitor software and processed with custom Python code [31] as described in [22]. 3D QPI was performed with Tomocube HT-X1 (Tomocube Inc., Daejeon, Korea). Refractive index maps *n*(*x, y, z*) from the Tomocube software were similarly analysed with our Python pipeline [22, 31].

Both 2D and 3D QPI rely on the linear relationship between RI and dry mass concentration (eq. 1). Apart from this, the two modalities use different light paths, reconstruction algorithms, assumptions, and post-processing steps in order to compute cell refractive index and height. None of the instruments was calibrated prior to the measurements, therefore the two modalities were used to cross-calibrate each other. The calibration procedure is described in Appendices B and C. In 3D QPI, voxel-wise RI maps are reconstructed tomographically, and cell height is then obtained directly from the 3D segmentation, which depends on the axial resolution (point-spread function). In 2D QPI, the measured phase (optical path length) is converted to an effective cell height by dividing it by the RI model from Appendix C.

### C. Cell culturing

Madin–Darby canine kidney cells, MDCK parental line (ATCC NBL-2 CCL-34, purchased in 2004) were cultured in complete Dulbecco’s Modified Eagle Medium (DMEM, Sigma-Aldrich). All cells were grown in the presence of 10 % FBS (Sigma-Aldrich) and 1 % penicillin/streptomycin (Sigma-Aldrich) at 37 ◦C in a humidified atmosphere containing 5 % CO_2_. The experiments were carried out at passaging numbers 2-20. Cells were seeded at different densities in 35 mm glass bottom dishes (MatTek, part no: P35G-1.5-14-C) with and without circular fibronectin patches 24-72 hours before imaging was started.

### D. Image acquisition

All data is from 14 different experiments and 6 biological replicates (different passaging numbers), derived from three separate frozen stocks of the same cell line. Each experiment produced a timelapse dataset of height *h*(*x, y, t*) over a field of view of approximately 567 × 567 µm^2^, with a duration of 10 − 15 hours. The lateral resolutions are Δ*x* = 0.5537 µm for Holomonitor and Δ*x* = 0.1554 µm for Tomocube. With our imaging conditions, we obtain temporal resolutions of Δ*t* = 5 min and Δ*t* = 15 min for Holomonitor and Tomocube, respectively.

### E. Image analysis

The cell detection, segmentation and tracking methods are described in detail in [22]. From this we obtain data of position **r**_*i*_(*t*), projected area *A*_*i*_(*t*), mean height 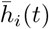, volume *V*_*i*_(*t*), and dry mass *m*_*i*_(*t*) of a segmented cell *i* at time-point *t*. Cell tracks are terminated 30 minutes before division events to exclude mitotic cells that round up, shed mass and divide. Mother and daughter cells are always treated as distinct cells with separate trajectories.

We quantify the spatial and temporal height variations in the monolayer as the relative standard deviations:

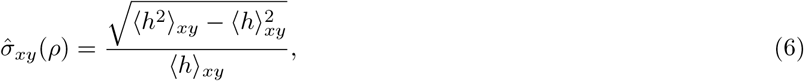

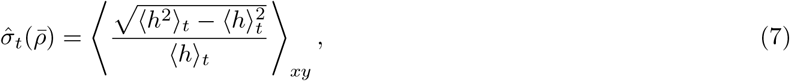

where *h* = *h*(*x, y, ρ*(*t*)). The temporal average is taken over 4 h, which has been found to be the timescale of oscillations in the projected area of MDCK [8]. 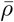 denotes the mean cell density in this time interval.

We use PIVlab [32] to obtain the velocity field of the monolayer. This is performed on Tomocube *n*(*x, y, t*) datasets that have higher lateral resolution and local contrasts. We use two multi-pass window sizes (40 µm and 20 µm) corresponding to 2 to 1 cell diameters, respectively, where the largest is chosen for robustness and the smallest to capture the displacements on the order of a cell size. The windows are moved with a step size of 10 µm. For frames of 3648 × 3648 pixels, we then obtain a 56 × 56 blockpixel velocity field. We remove spurious velocity vectors and replace them with an interpolation of the remaining vector field. Spurious vectors are defined as any vector with magnitude greater than 40 µm/h.

We compare the mass flux and the mass change in the monolayer. We use finite volume to compute the mass flux in and out of every pixel of the velocity field. First, we compute the mass field *m*(*x, y, t*) = *α*^−1^*h*(*x, y, t*) · Δ*n*(*x, y, t*). We downsample it to the same resolution as the velocity by excluding 32 pixels along the edge, then splitting it into 10 × 10 µm^2^ blocks and replacing each block with the average value of its pixels. Then, we shift both velocity and mass fields onto the edges of every blockpixel and compute the mass flux as ∑_*i*_ *m*_*i*_ · *v*_*i*_, where *i* runs over the four faces of each blockpixel. We compute the mass change as the difference between consecutive frames of the downsampled mass field. We also calculate the space-time autocorrelation functions *C*_*X*_ (**r**, *t*) = *δX*(**r**^′^, *t*^′^) *δX*(**r**^′^ + **r**, *t*^′^ + *t*) _**r***′,t′*_, where *δX*(**r**, *t*) = *X*(**r**, *t*) − ⟨*X*(**r**, *t*)⟩ of *X* ∈ {∂_*t*_*m*, ∇ · (*m***v**)}.

## III. RESULTS

All the results presented here, except for refractive index distributions and velocity fields, are derived from data from both 2D and 3D QPI measurements.

### A. Dry mass concentration varies weakly across a monolayer

There are large differences in the refractive index inside a cell depending on the local concentration of proteins, lipids, etc. (see Figure 1A), and therefore the intracellular dry mass concentration is not uniform. We tested whether the *mean* concentration of cellular dry mass varies significantly in the monolayer. Figure 1B shows the spatial variation of a dry mass concentration field *c*_*d*_(*r*; *x, y*) in a confluent monolayer Gaussian filtered with radius *r* = 0.9 × cell radius [22]. On this coarse-grained level we observe spatial structure, and the most extreme regions deviate from the mean by up to 20 %. However, the distribution of *c*_*d*_(*r*; *x, y*) (Figure 1C) is relatively narrow, with a standard deviation (SD) of ≈ 4.5 %. This SD is three times higher than for three other mammalian cell lines [33] and half the reported value for MDCK-II, HeLa and NIH3T3 in [20]. Since cell height is inversely proportional to dry mass concentration (eq. (3)), the 4.5% fluctuations add to the 3% precision in phase determination [22] and our height determination has a precision of 5.4%.

**FIG. 1.**
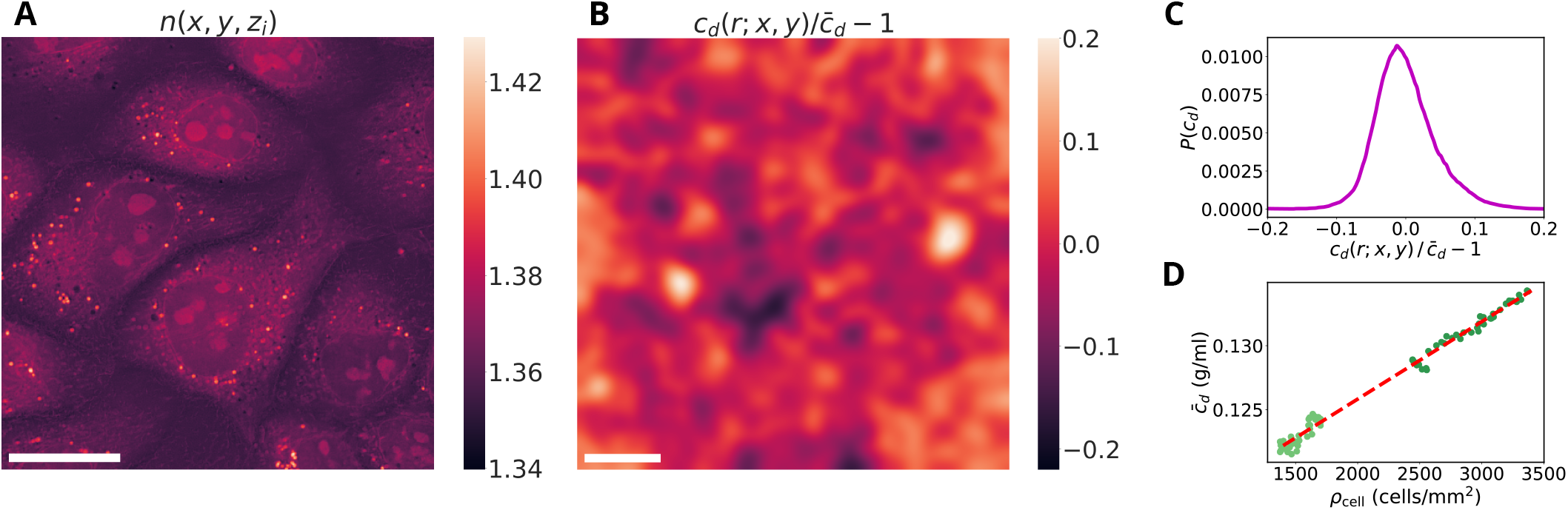
Dry mass concentration varies weakly across a monolayer. **A:** Refractive index *n*(*x, y, z*_*i*_) in a single optical section at *z*_*i*_ = (0.8 ± 0.3) µm. Scale bar: 20 µm. **B:** Spatial variation of the coarse-grained dry mass concentration field *c*_*d*_(*r*; *x, y*) across a confluent monolayer. Colours indicate deviations from the mean, 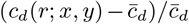. Scale bar: 100 µm. **C:** Probability density function of *c*_*d*_(*r*; *x, y*) plotted as deviation from the mean. The standard deviation is ≈ 4.5 %. **D:** Dry mass concentration as function of cell density. The dry mass concentration varies *c*_*d*_*/ρ* = (6.06 ± 0.08) 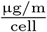 and *c*_*d*_ = 0.130 g/ml at *ρ* = 2700 cells/mm^2^

We also plot the mean dry mass concentration in the monolayer as function of cell density in Fig. 1D and see that it increases linearly with cell density as *c*_*d*_ = 0.113 + (6.06 ± 0.08)*ρ*_cell_ µg/ml. This is slightly higher than previous estimates for MDCK based on Raman and SRS measurements combined with 3D QPI [34] (see Table I). Note that Fig. 1D contrasts with the erroneous Figure 6 in [22] (see Appendix C).

**TABLE I.** Mean refractive index and dry mass concentration of MDCK from literature. The values in parenthesis have been derived from the other column using eq. (1), with *n*_0_ = 1.337.

| $\bar{n}$ | $c_d$ [g/ml] | Notes |
| --- | --- | --- |
| 1.399 | (0.326) | Single-cell RI from an on-chip Fabry–Pérot cavity resonator [59]. |
| $1.383 \pm 0.001$ | $(0.242 \pm 0.005)$ | Single-cell RI from a grating resonant cavity with optical trapping [60]. |
| 1.381 | (0.231) | Single-cell RI from a microfluidic cell-culture chip [61]. |
| $(1.355 \pm 0.002)$ | $0.095 \pm 0.11$ | Normalized stimulated Raman scattering imaging (NoRI)[24]. |
| $(1.357\text{--}1.365)$ | $0.105\text{--}0.147$ | Same NoRI method as [24] on confluent monolayers at $\rho_{\text{cell}} = 4000\text{--}8000$ cells/mm<sup>2</sup> [34]. |
| $1.363\text{--}1.368$ | $(0.138\text{--}0.164)$ | Our estimation at cell density $\rho_{\text{cell}} = 4000\text{--}8000$ cells/mm<sup>2</sup>. |

In summary, dry mass concentration increases with cell density and exhibits small but measurable spatial and temporal fluctuations. The dry mass-based phase signal is therefore proportional to cell height *h*(*x, y*) with a precision of ∼ 5%. This justifies treating 2D QPI height measurements and 3D QPI height measurements as probes of the same quantity, and we use height data from both modalities in the following analysis of epithelial height dynamics.

### B. Height and volume distributions depend on cell density

We observe a substantial height variation in MDCK monolayers, as can be seen in Figure 2A. The global cell number density in the Field Of View (FOV) is *ρ*_cell_ = 1650 cells/mm^2^, but local cell density varies considerably. Cells in denser regions occupy a smaller area and are taller than cells in less dense regions. We also observe a few very tall cells (almost twice the mean monolayer height) that are rounding up as mitosis approaches. Within a cell, the cell centre tends to be taller than at the cell–cell contacts.

**FIG. 2.**
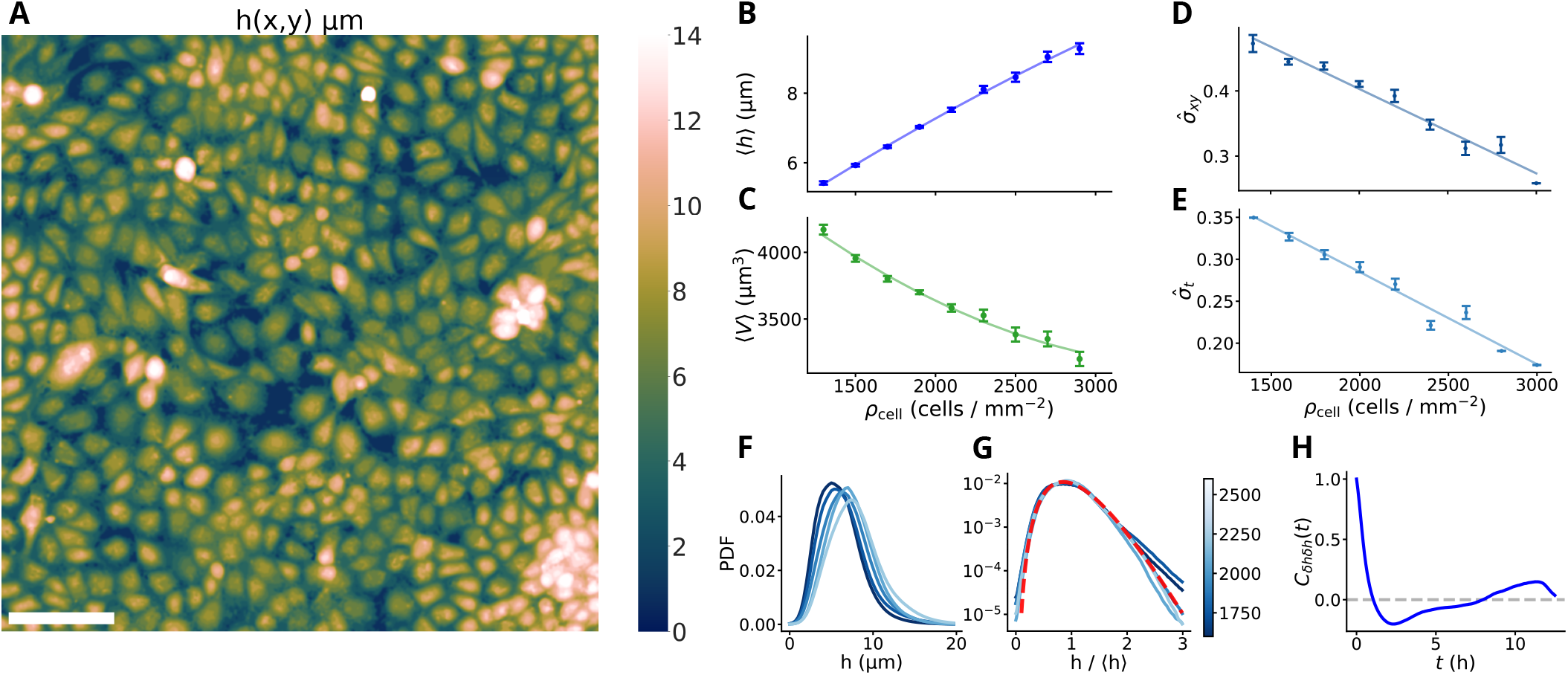
MDCK height fields are heterogeneous and dynamic. **A:** Heatmap of the height field of an MDCK monolayer at cell density 1650 cells/mm^2^. The scalebar represents 100 µm. The monolayer is confluent although in some sparse region the height is very low. **B:** Mean monolayer height as function of cell density *ρ*_cell_ based on all 2D and 3D QPI measurements. A weighted second order polynomial is fitted to the data: 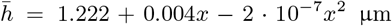. **C:** Mean cell volume ⟨*V* ⟩_*xy*_ = ⟨*h*⟩_*xy*_*/ρ*_cell_ as function of cell density *ρ*_cell_. Weighted second order polynomial fit: 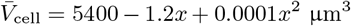. **D:** Spatial height variations: Relative standard deviation, 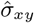 in space over monolayers as function of cell density. Weighted linear fit: 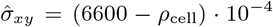. **E:** Temporal height variations: Relative standard deviation, 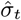 in time at the same position in space as function of cell density. The average is taken over a time interval of 4 h and is plotted against the average cell density during this interval. Weighted linear fit: 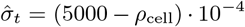. **B-E:** Data for these plots was averaged over 14 experiments and 6 biological replicates on glass and fibronectin using two QPI modalities. The errorbars represent the standard error of the mean across all biological replicates. **F:** Height distributions at different cell densities. The cell densities vary from 1600 − 2600 mm^−2^, with dark lines denoting low densities and bright lines denoting high densities. **G:** Logarithmic probability density function of *h/*⟨*h*⟩_*xy*_ for the same cell densities as F. A gamma distribution is fitted to the data (red dashed line), with the shape parameter *α* = 5.9 and the scale parameter *θ* = 0.17. **H:** Mean detrended height autocorrelation of *h*(*x, y*) of each pixel, where the height of every pixel has been detrended to remove the height increase due to cell growth. The mean is taken over all initial cell number densities. **F-H:** from *one* experiment on glass. All data in **B-G** are distributions and averages of pixel-wise heights as displayed in A.

Figures 2B-E show the aggregate results of all 14 data sets for confluent monolayers at cell densities *ρ*_cell_ = 1300 − 3000 cells/mm^2^. The mean height of the monolayer shows an approximately linear increase from 5.5 µm to 9 µm (Figure 2B), which means that cells elongate in the *z*-direction as cell density increases and the mean area per cell, ⟨*A*⟩_*xy*_ = 1*/ρ*_cell_, decreases. Although the mean height increases with cell density, Figure 2C shows that also the mean cell volume, ⟨*V* ⟩_*xy*_ = ⟨*h*⟩_*xy*_*/ρ*_cell_, decreases with cell density. This is not due to contact inhibition of proliferation, which is the most commonly discussed mechanical effect of confinement on tissue growth [35, 36], but rather due to contact inhibition of cell *size*, which is also consistent with the results of Devany et al. [18]. However, their height measurements by confocal fluorescence microscopy did not resolve any increase in height with increasing cell density. This led them to the conclusion that cells divide but do not grow, which is not supported by our data.

As seen in Figure 2D, there are large spatial height variations 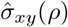 throughout the monolayer. We see that the height variations are largest at low cell density, and fall from 48% to 28% at high density, where the monolayer becomes more homogeneous. The height at any pixel in the FOV varies in time as well, which is shown in Figure 2E. Like in the case of spatial height fluctuations, the temporal height fluctuations 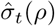 are largest at low cell density and drop from 35% to 18% as the cell density increases.

The height probability density functions P(*h*(*x, y, ρ*_cell_)) from a single experiment are shown in Figure 2F. The cell density ranges from 1600 cells/mm^2^ to 2600 cells/mm^2^. As the cell density increases, both the mean and the width of the height distribution increase. To compare distributions across densities, we rescale the height by its mean and plot the corresponding distributions in Figure 2G. This rescaling produces a partial data collapse, and at higher densities the rescaled distributions are well described by a gamma-like form. The tails of the distribution are suppressed when density increases, which results in the reduced height variation at higher density shown in Figure 2D,E.

Some of the height variation in Figure 2D,E come from intracellular variation, since each cell is the highest at their centre and the lowest at their periphery. A Gaussian-filtered height field [22] (see Figure 6) that represents average cell heights shows spatial variation of 30%. This spatial height variation is six times larger than those found by Zehnder et al. [8] (±4.7 % height variation in space at cell density 1300-1400 mm^−2^) using confocal fluorescence microscopy. Thus, our results do not qualify the MDCK layers as “flat” as described by Zehnder et al. [8], but rather resemble the qualitative description of height fluctuations by Thiagarajan et al [10, 37].

We have shown that cell heights in MDCK monolayers display dynamic spatial heterogeneity, with height distributions well described by a gamma distribution that persists even at high densities. The timescale of these fluctuations can be found using the height autocorrelation function *C*_*δhδh*_. The result is shown in Figure 2H. *C*_*δhδh*_ has a minimum at around 2.5 h, similar to what has been observed for the autocorrelation of cell areas [8]. Comparing it to the Gaussian-filtered autocorrelation in Figure 6G, we see that the negative peak becomes more pronounced when we average it over the radius of a cell, indicating that cell heights oscillate with a period of approximately 5 h.

### C. Cell-resolved dynamics: apparent mass loss

We have found that cells regulate the dry mass concentration and therefore the dry mass and volume of the cells are proportional to first approximation. We have also shown that cell heights fluctuate significantly over time. One question that arises is: are the cell area and cell height directly coupled through a constant cell volume? In order to understand how the three are dynamically linked, we measure the mean height, area, and volume of segmented cells in the 2.5D approximation (see eqs. (4) and (5)) and follow the evolution of individual cells in time. During the cell cycle, a cell will grow before eventually dividing, and thus, the cell volume is not expected to stay conserved on the length scale of a cell cycle. The question is rather whether the cell volume is strictly determined by the cell cycle or if it is affected by fluctuations in cell height and density.

Figure 3A shows the inverse area and height of nine representative cells. If the cell volumes were constant, the height and inverse area of a cell would be proportional. Although the height and inverse area curves show a large degree of correlation, we observe that there is a certain degree of deviation between the two. To better quantify this, the correlation between height *h*_*i*_*/*⟨*h*⟩_*i*_ and area *A*_*i*_*/*⟨*A*⟩_*i*_ for 5400 cells in 16 experiments is shown in Figure 3D. The Pearson correlation coefficient is 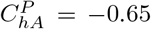, showing that height and area are only partially correlated during the collective dynamics of the cell sheet. This clearly signifies that the measured cell volumes are not constant during dynamic variation of the cell area, as is also evident from the volume tracks in Figure 3B. However, the dry mass concentration of individual cells in the same figure is remarkably constant with a standard deviation of only 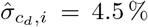. If the dry mass concentration is regulated and the 2.5D approximation is correct, the periods of decreasing volume observed in Figure 3B signify that cells periodically shed up to 10-15 % of their dry mass over periods of 1-2 hours. Because the periods surrounding mitosis are excluded from these tracks, this apparent mass loss is a feature of non-mitotic cells and cannot be attributed to the dry-mass shedding that accompanies mitosis [33].

**FIG. 3.**
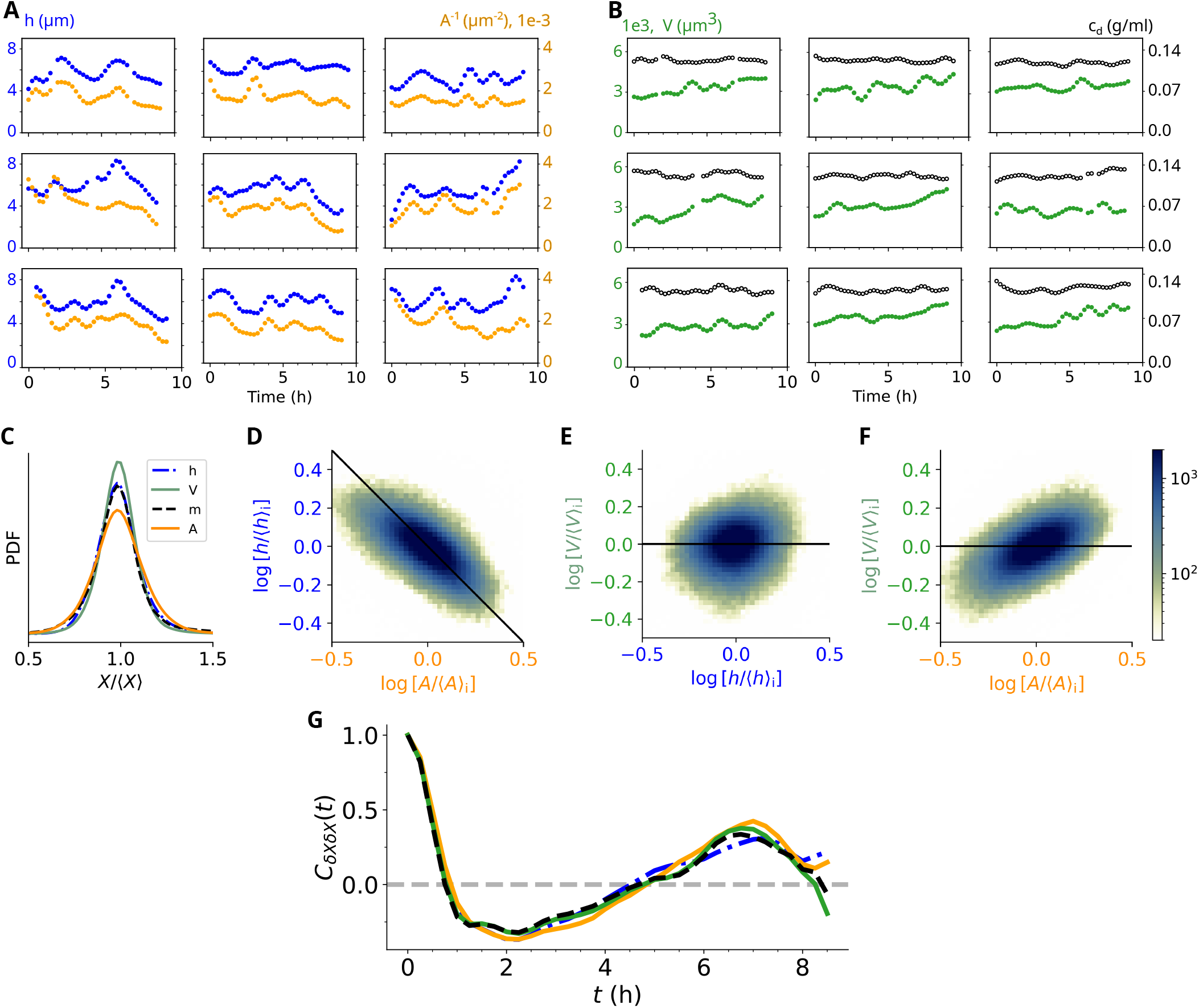
Cell height, projected area and volume fluctuate synchronously. **A:** Representative examples of height (blue dots) and inverse area (orange dots) of tracked single cells in monolayers of cell density *ρ*_cell_ = 1300 − 1700 cell/mm^2^. **B:** Volume (green dots) and dry mass concentration (black, open dots) of the same tracked cells as in A. **C:** Probability density functions of rescaled detrended cell height (h), area (A), volume (V), and dry mass (m). **D:** Scatter plot of individual detrended cell height vs. area. **E:** Scatter plot of individual detrended cell volume vs. height. **F:** Scatter plot of individual detrended cell volume vs. area. **C-F** contains data from 16 experiments at cell density *ρ*_cell_ = 1300 − 3300 cell/mm^2^. ⟨ · ⟩_*i*_ denote the average over 5400 cell tracks. The black lines in **D-F** are the expected relation for 2.5D/prismatic cells with constant volume. **G:** Time correlation functions of detrended height (blue, dash-dotted line), area (orange, solid line), volume (green, solid line) and dry mass (black, dashed line) of two experiments at all cell densities.

In Figure 3E,F we plot height and area against volume and see that the cells do not conserve volume when area and height fluctuates (black horizontal line). The correlation is much stronger between area and volume 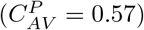 than between height and volume 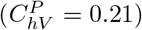. Thus, volume fluctuations seem to be mainly driven by area fluctuations, with minor contribution from height fluctuations. As shown in Figure 3G, the time autocorrelation functions of area, height, and volume (dry mass) overlap, suggesting that they are dynamically coupled during the collective motion of the monolayer. This also implies periodic loss of cellular dry mass, and/or non-prismatic cell geometry. The periodic loss of cellular dry mass is a phenomenon that, to our knowledge, has not been reported before. In a biological context, it is counter-intuitive that cells would actively discard biomass, given its energetic cost, unless this confers an advantage at the population level.

We emphasise that the cell volume we measure is the volume under the segmented apical surface. This assumes the cell-cell junctions to be vertical, thus treating cells as prisms. However, confocal images of vertical sections through epithelial monolayers show that the lateral membranes are not always vertical [38]. More complex cell geometries have been shown to be necessary in curved epithelia [11, 39]. From a geometric perspective, the natural extensions beyond prisms are frustums, prismatoids, and scutoids, none of which need to have equal basal and apical areas. An illustration of an epithelial layer with non-vertical lateral membranes and three space-filling frustums is shown in Figure 4A. We tested whether non-prismatic cell shapes could produce the apparent volume fluctuations of Figure 3 even when the true volume is constant (see the model description in Appendix A 4).

**FIG. 4.**
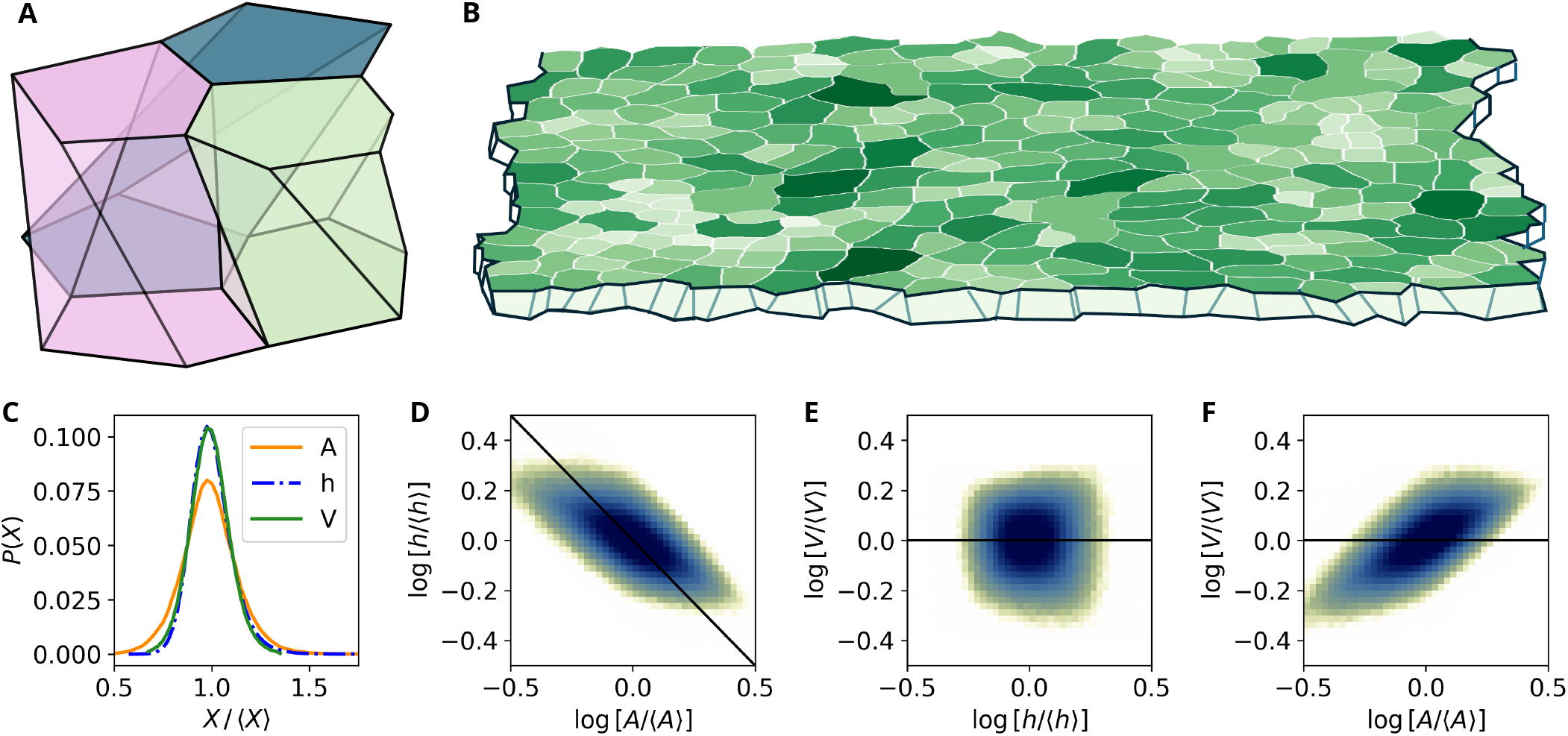
Frustum-based cell models reproduce experimental height–area–volume correlations. **A:** Example of three frustum cells in contact: apical and basal areas are parallel but can have different size and position of the polygon. **B:** Measured areas of MDCK layer with non-vertical lateral cell membranes sketched to indicate a confluent layer of frustums. **C:** Probability density functions (PDFs) of the height, area, and projected volume (height times apical area) of frustums with constant volume. *A/*⟨*A*⟩ is taken from the experimental area distribution (Figure 3C). Apical and basal areas are independently drawn from *A/*⟨*A*⟩ and are then used to calculate height and projected volume according to eqs. (A3-A2). **D-F:** Density plots relating height, projected (apical) area and volume (height times apical area) for frustums with constant volume. The colorbar is the same as in Fig. 3. The black lines are relations for prisms.

The height and projected volume distributions calculated using our model for the frustum case are shown in Figure 4C, and the three pairwise correlations between projected area, height and measured projected volume are shown in Figure 4D-F. Qualitatively, these correlations closely resemble the experimentally observed correlations in Figure 3D-F. The area is strongly correlated with both height and volume, whereas height and volume themselves are uncorrelated. Thus, a frustum-shaped cell with constant true volume would yield fluctuating projected volumes and reproduce behaviour very similar to what is observed experimentally, even though the actual cell volume might not change. The corresponding results for prismatoid cells are shown in Figure 9. Because the result is sensitive to correlations in the model setup, we find it unrealistic that the frustum geometry can account for all observed variation while maintaining reasonable correlations between apical and basal areas (see Section A 4).

### D. Continuum distributions and motion: continuity fails at cellular scale

We have shown that the projected volume/dry mass of segmented cells is not constant and that segmented cells experience periods of apparent mass loss. For a complementary and more robust analysis, we now consider conservation in a continuous mass field *m*(*x, y, t*).

In Appendix A 3 we compute the structure factor *S*(**q**, *ω*) which is the Fourier transform in space and time of the mass field *m*(*x, y, t*) (Figure 7). The static structure factor crosses over from *S*_*m*_(*q*) ∼ *q*^−2^ at long wavelengths, the signature of scale-free density fluctuations, to *q*^−4^ at the cell scale. The dynamic structure factor reveals a Brillouin-like propagating mode whose frequency and damping both scale linearly with *q*, corresponding to an elastic wave with group velocity *c*_*s*_ ≃ 15 µm/h. The same crossover and propagating mode appear across three independent imaging modalities, phase contrast [40], fluorescence intensity [9], and the direct dry-mass field measured here. This validates our mass field and shows that the monolayer supports propagating elastic excitations rather than purely diffusive relaxation. Having thus validated the mass field, we now return to the question of mass conservation.

The continuum version of the 2.5D approximation is plug-flow in a thin film. We consider the fluid to be incompressible, since our measurements show that the dry mass concentration varies only a few percent under our conditions. The height-averaged mass continuity equation for a thin plug-flow film is

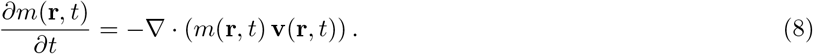

Note that since cells are virtually incompressible, the mass *m*(**r**, *t*) may be exchanged with the height *h*(**r**, *t*) in the equation above.

In Figures 5A,B we show the instantaneous mass change ∂_*t*_*m* and the divergence of mass flux ∇·(*m***v**) in a monolayer of cell density *ρ*_cell_ = 1700 cells/mm^−2^. In the supplementary movie S4 we display these two images side by side for every time point of the entire experiment, where we have smoothed both fields with a Gaussian blur with kernel radius *r* = 30 × 30 µm^2^ to capture the collective movement rather than the instantaneous displacement of single cells.

**FIG. 5.**
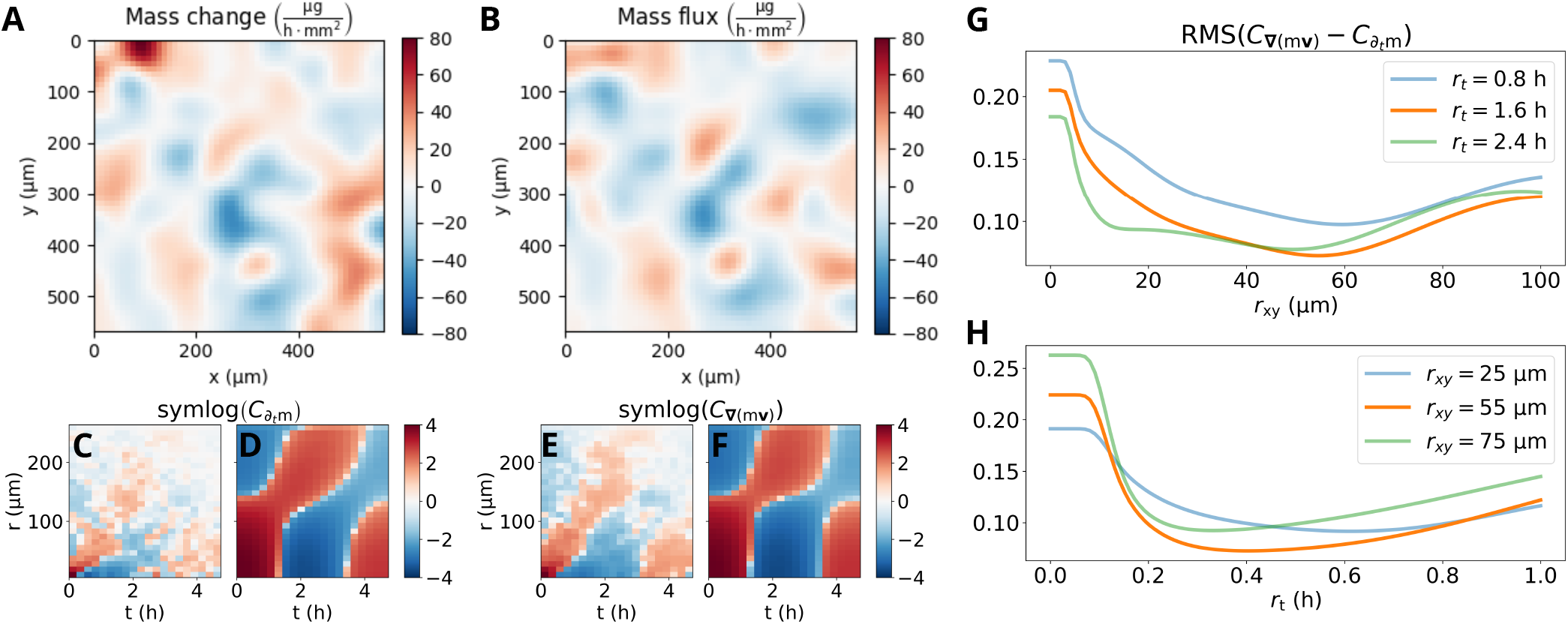
The monolayer mass field only obeys mass conservation on longer time and length scales. **A:** Instantaneous mass change, *dm/dt*, between two frames, *dt* = 0.25 h, in monolayer of cell density *ρ*_cell_ = 1700 cells/mm^2^. **B:** Instantaneous mass flux, −∇ · (*m* **v**), at the same instance. **A, B:** Mass conservation by the continuity equation requires mass change and mass flux to be equal. Both *dm/dt* and ∇·(*m***v**) are computed as averages of the flux/change within a block of 10 × 10 µm^2^, then a Gaussian filter with kernel radius 30 × 30 µm^2^ is applied to each frame. **C, E:** Space-time autocorrelation functions *C*_*X*_ (**r**, *t*) are evaluated as for A and B. The result is displayed as a symlog, that is, a logarithmic colour scale applied to both positive and negative numbers. Mass conservation by the continuity equation requires C and E to be equal. C and E are averaged over all instances and are therefore a better comparison than A and B. **D, F:** Same as C and E, but the data has been Gaussian filtered with kernel radius of (*r*_*xy*_, *r*_*t*_) = (55 µm, 1.6 h) *before* calculating the correlation function. **G, H:** Root-mean-square of the residuals of the space-time autocorrelation functions as function of spatial (**G**) and temporal (**H**) smoothing parameters. The kernel size (*r*_*t*_ = 1.6 h, *r*_*xy*_ = 55 µm) is found to be the minimum.

**FIG. 6.**
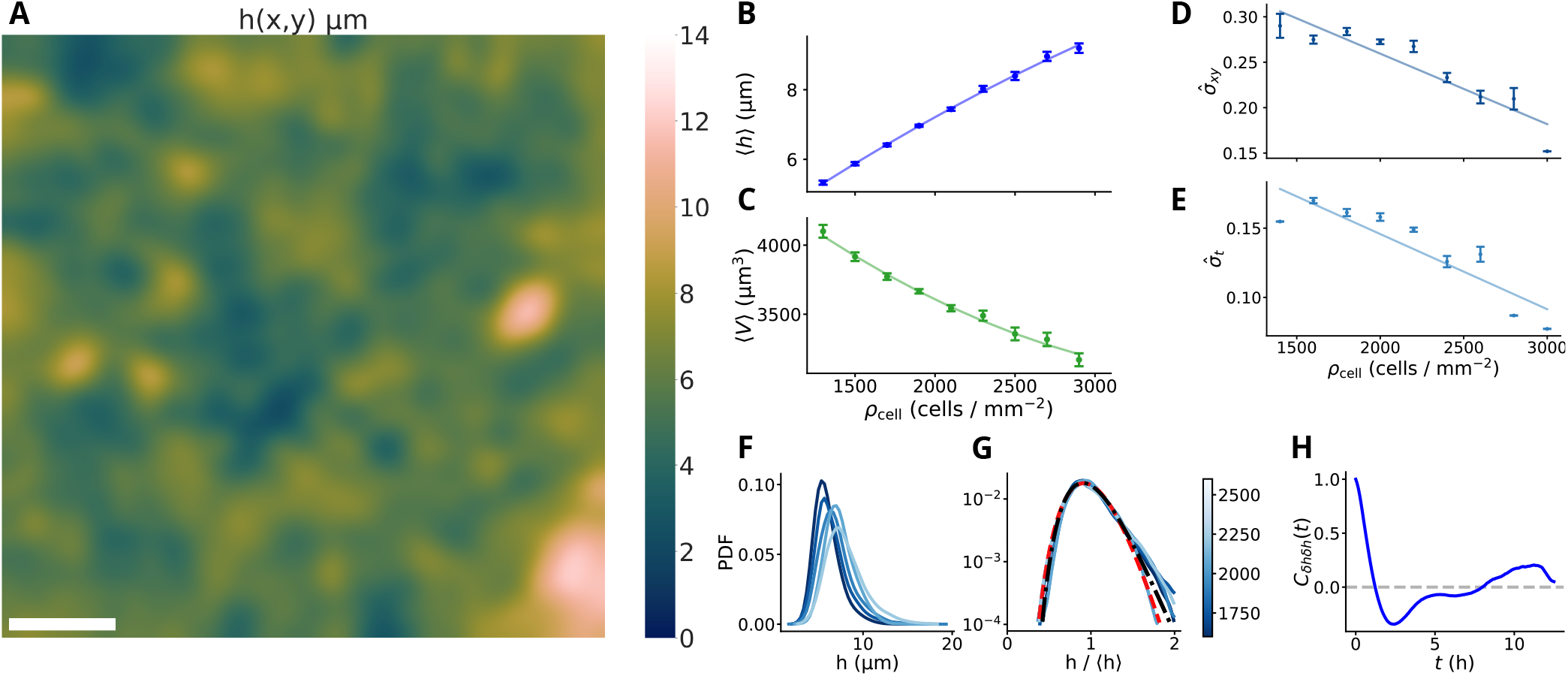
Characteristic Gaussian filtered height distributions of MDCK monolayers. All data are disc-wise averages that represent average cell heights as shown previously [22]. The original data is exactly the same as in Figure 2 but disc-wise averaged to remove the height variation across each cell and therefore represent distributions of average cell height. **A:** Image of MDCK monolayer where intensity equals height. The scalebar represents 100 µm. **B:** Mean monolayer height as function of cell density for 14 experiments and 6 biological replicates on glass and fibronectin using 2 different measurement techniques. A linear function is fitted to the data 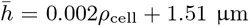. **C:** Mean monolayer volume as function of cell density for 14 experiments and 6 biological replicates on glass and fibronectin using 2 different measurement techniques. A linear function is fitted to the data 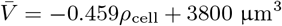. **D:** Spatial fluctuations: Relative SD, 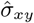 in space over monolayers as function of cell density. A linear function is fitted to the data 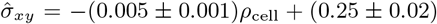. **E:** Temporal fluctuations: Relative SD, 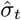 in time at the same position is space as function of cell density. 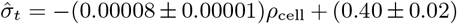. **F:** Height distributions at different cell densities for one experiment. The cell densities vary from 1400 − 2300 cells/mm^2^, with dark lines denoting low densities and bright lines denoting high densities. **G:** Logarithmic probability density function (PDF) of *h/*⟨*h*⟩ for the same cell number densities as in E. A gamma distribution (red dashed line, *α* = 16.74 ± 0.05, *θ* = 0.06, *χ*^2^ = 0.02) and a log-normal distribution (black dash-dotted line, µ=0.25, *σ* = 0.95, *χ*^2^ = 0.005) are fitted to the data. **H:** Mean height autocorrelation of detrended Gaussian filtered heights at fixed points in space. The mean is taken over all initial cell number densities larger than *ρ*_cell_ = 1600 cells/mm^2^.

**FIG. 7.**
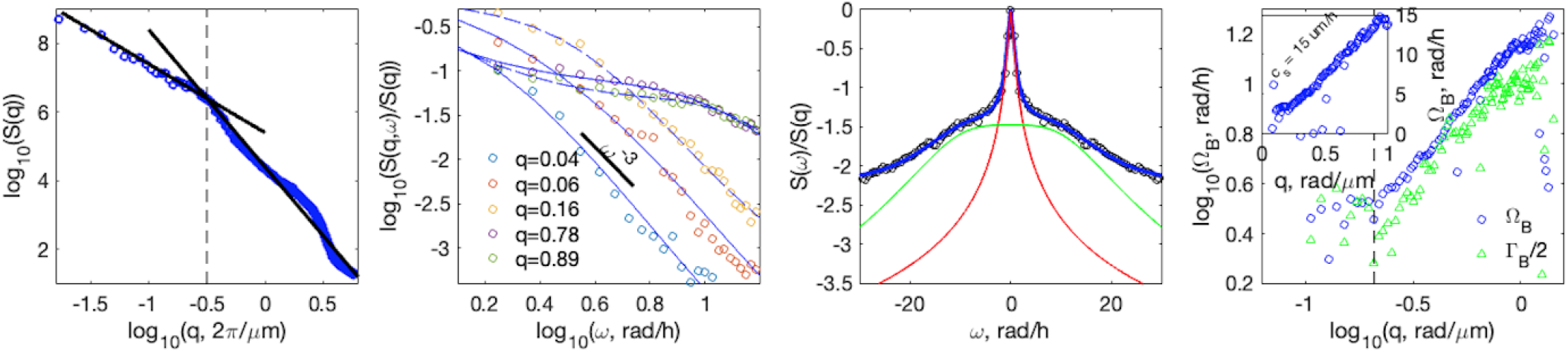
Structure factors of epithelial mass field. **A-C:** Properties of structure factors of monolayer mass *m*(**r**, *t*), *S*_*m*_(*q, ω*) = ⟨|ℱ(*m*(**r**, *t*))|^2^⟩_*ϕ*_ where ℱ(*m*) is the Fourier transform of *m* in spatial *q* and temporal *ω* frequencies for monolayer densities *ρ*_cell_ ∈ [1300, 2400] cells/mm^2^. **A:** The static structure factor *S*_*m*_(*q*, 0) ∼ *q*^*n*^ has a crossover from to *n* = −2 to *n* = −4 at 20 µm, corresponding approximately to a cell diameter. **B:** The dynamic structure factor *S*_*m*_(*q, ω*)*/S*_*m*_(*q*) for several *q*. For intercellular length-scales (2*π/q >* 20 µm) and large *ω* the structure factor scales as *S*_*m*_(*q, ω*) ∼ *ω*^−3^, which corresponds to sub-diffusive displacements. **C:** Example of fitting *S*_*m*_(*q, ω*) with a Lorentzian Rayleigh peak (red line) and a DHO type Brillouin peak (green line). **D:** The frequency Ω_*B*_ and half-width Γ_*B*_ */*2 of the Brillouin peak both scale linearly with *q*, corresponding to a speed of 15 µm/h.

**FIG. 8.**
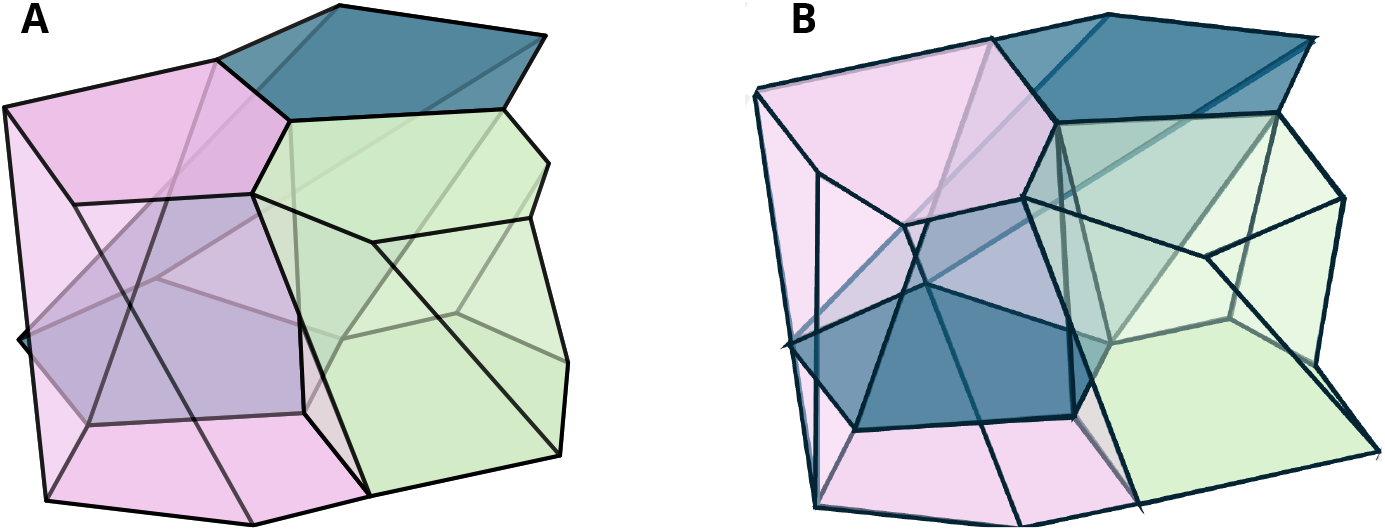
**A:** Example of three frustum cells in contact: apical and basal areas are parallel but can have different size and position of the polygon **B:** Example of three prismatoid cells in contact: apical and basal areas are parallel but can have different size, position and vertexes of the polygon

**FIG. 9.**
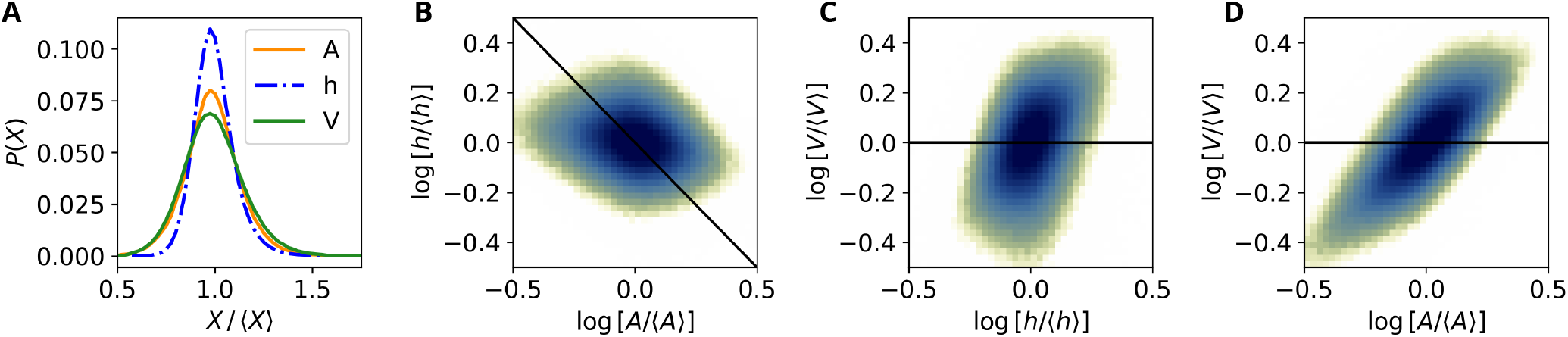
Distributions of height, projected area and projected volume of prismatoids. **A:** Probability density functions (PDFs) of the height, area, and projected volume (height times apical area) of prismatoids with constant volume. *A*_*a*_, *A*_*m*_, and *A*_*b*_ are drawn independently from the experimental area distribution *A/*⟨*A*⟩ and are used to calculate *h* and *V*_*m*_ according to eqs. (A6-A5). **B-D:** Density plots relating height, projected area and projected volume (height times apical area) for prismatoids with constant volume. The Pearson correlation coefficients are: 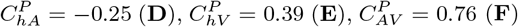. The colorbar is the same as in Fig. 3. The black lines are relations for prisms.

The lack of agreement between the two fields in Figures 5A,B on length scales below the size of 2 cells indicates that the continuity equation is not satisfied instantaneously at this resolution. This implies either that mass is not strictly conserved at this scale or that the flow is not a plug flow. This is consistent with our conclusion from analysis based on cell segmentation.

To obtain an averaged description in space and time, we compute the space–time autocorrelation functions of ∂_*t*_*m* and ∇ · (*m***v**), shown in Figure 5C, E. We observe that there is some qualitative similarity between the two, but the quantitative agreement is best when we apply a 2D Gaussian filter with kernel radii *r*_*xy*_ = 55 µm and *r*_*t*_ = 1.6 h as shown in Figure 5D and F. We vary the spatial and temporal kernel radius, and compute the root-mean-square of the correlations in Fig. 5G,H. Thus, the continuity equation and mass conservation appear to hold at these larger spatio-temporal scales, but not at the smallest scales resolved in our measurements.

## IV. DISCUSSION

### A. Dry mass concentration is maintained within a narrow range

There is recent compelling evidence of strong regulation of dry mass concentration [20, 24, 41, 42]. The exact mechanisms of regulation of dry mass concentration are still unknown, but more theories are forming that link elevated mass concentration to decreased protein synthesis rate [41] and suggest that volume regulation and protein biosynthesis are controlled by two independent mechanisms [42].

Our measurements add to this picture by showing that, even in dynamically pulsating and collectively migrating MDCK monolayers, the coarse-grained dry mass concentration varies only by a few percent across space and time, whereas cell height exhibits much larger fluctuations. This comparatively small variation in dry mass concentration places constraints on the physical mechanisms that can drive height and area fluctuations. In particular, it rules out intercellular fluid transport as the dominant driver of epithelial height and area dynamics, as previously proposed [8, 9, 43]. In those studies the pulsations are attributed to the flow of water between cells — through gap junctions or in exchange with the medium — driven by osmotic pressure gradients. Such transport carries water and small ions but not the macromolecular dry mass, which cannot cross the gap junctions in appreciable amounts. Therefore, a cell expanding by water uptake would be diluted and a contracting cell would be concentrated. Water-driven volume fluctuations of the observed amplitude (tens of percent) would thus produce dry mass concentration fluctuations of comparable size, which the few-percent constancy of *c*_*d*_ excludes.

Since common belief until 2022 was that mass regulation and water volume regulation in cells were independent, one may indeed reinterpret earlier results on volume fluctuations. It is clear that the fast cellular response to osmolarity changes is due to the regulation of osmotic pressure and can be explained by the pump-leak model [17]. However, due to this model and the inability of commonly used measurement techniques to measure the dry mass of the cells, all volume reductions were interpreted as a result of water efflux [24]. In view of our current knowledge, one should examine whether volume reducing cells actually shed, excrete, or exchange mass as they were found to do during mitosis [33].

Consistent with this view, Chmiel and Gardel [44] found that confluent MDCK monolayers, unlike isolated cells, fail to restore their volume after a hyperosmotic shock, single-cell NKCC-mediated volume homeostasis being suppressed by the tight junctions of mature epithelium. Their work concerns water-driven volume changes under imposed osmotic stress rather than dry mass concentration during spontaneous pulsations, but it reinforces that volume regulation in confluent tissue differs qualitatively from the single-cell picture and should be reinterpreted alongside direct dry mass measurements such as ours.

### B. Mass conservation and cell volume and shape changes

We have shown that measures based on both cell segmentation and on continuum fields show a lack of volume or mass conservation on the cellular scale. We present two main interpretations of our observations of volume/mass fluctuations in synchronisation with the large scale density fluctuations of the monolayer: a) cells periodically excrete mass, or b) cells twist, turn, and do not behave like “prisms”. The two interpretations are not mutually exclusive; that is, both can be true.

#### a. Monolayer dynamics compress and stretch cells that respond by varying mass

The idea of mechanical feedback on tissue growth is well established [35, 36, 45–47]. Stretched epithelia grow faster than compressed epithelia. Although there is most experimental evidence that mechanical stress affects cell proliferation rate, there is growing evidence that metabolic rates adapt to mechanical stress [48]. An increase in osmotic pressure reduces cell size, ECM stiffness has been reported to result in varying effects on cell size depending on cell type, and shear stress decreases cell size [48]. Confinement has also been shown to affect cell size [18]. Flow-induced shear stress of 0.2 Pa on MDCK monolayers has been shown to reduce volume by up to 25% in less than 1 hour [49]. Volume reduction was hypothesised to be due to water efflux, but recent evidence of strong regulation of mass concentration [20, 24, 41, 42] and our results that mass concentration is only weakly varying even during large collective motion suggests that the volume reduction response is due to mass reduction of MDCK cells.

The timescale of mass variation we find is in good agreement with Liu et al. [50] that showed growth rate variations with an average 5 h period of mass growth oscillations in HeLa cells. The growth rates they measured were always positive, that is, single HeLa cells do not shed mass. However, there is no reason to expect identical behaviour in the HeLa cancer cell line and the MDCK cell line that is known to deposit well organised basal lamina [51].

Do cells lose dry mass? Few studies have used techniques that can answer this question. Miettinen et al. [33] recently used a suspended microchannel resonator (SMR) to measure the dry mass of single cells and found that during mitosis they exocytose a significant fraction of their biomass. Since our tracked-cell analysis excludes periods surrounding mitosis, the apparent mass loss we observe occurs in interphase cells and is therefore distinct from this mitotic exocytosis. Renal epithelia like MDCK excrete laminin, fibronectin, collagen, proteoglycans, and nidogen to build their own basement layer [51]. To our knowledge, it is not known whether the exocytocis of the extracellular matrix is continuous or occurs in bursts for immature epithelial monolayers (for cell densities, 1300 − 3000 cells/mm^2^, which we have studied here [38]). Thus, it is possible that the excretion of ECM is pulsatile and synchronised with the collective motion of the monolayer.

#### b. Frustum-shaped cells affect projected measures of cell mass

Almost all analysis of flat epithelial monolayers is based on the assumption that cells can be modelled as prisms: the apical and basal surfaces have the same projected area and the lateral sides are vertical [6]. This is also the basis for 2D models and several 3D models of epithelia. Non-prismatic cell shapes have so far been introduced almost exclusively in the context of tissue curvature, where they are a geometric necessity: frustums describe the “bottle” cells of invaginating epithelia [5, 13], and scutoids resolve the apico-basal mismatch of neighbour relations in bent or tubular epithelia [11, 12]. Hannezo et al. [5] model planar cells as hexagonal prisms and introduce apical–basal area asymmetry only once the sheet bends. Recent 3D segmentation of curved epithelial showed scutoid and skewed cell shapes [39]. However, Dawney et al. [38] show images of cross-sections of flat, mature MDCK layers that indicate that the lateral cell membranes are not always vertical.

Our analysis shows that the prismatic assumption is inadequate even for flat monolayers. The similarity of the area, height and volume correlations from experiments (Fig. 3D-F) and from the frustum model (Fig. 4D-F) suggest that the frustum shape deformations of the cells can explain a large portion of our data. This is consistent with the direct confocal cross-sections of flat MDCK layers reported by Dawney et al. [38]. However, there are some limiting factors: 1) our monolayers are “squamous”, the range of average aspect ratios in our experiments is only 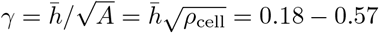. The frustum model can reproduce the experimental data only if a small fraction of the cells have large differences in the apical and basal area. 2) The height–volume correlation of the frustum model yields 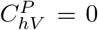 by construction and the prismatoid model yields 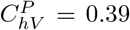, while the experimental value is 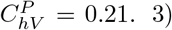) If we draw the apical area from the experimental distribution and the ratio between apical and basal area from a Gaussian distribution, the mentioned correlations become completely different. We conclude that non-prismatic geometry can account for most of the apparent volume fluctuations, but we are not convinced that it can account for their full amplitude.

#### c. Weighing the two interpretations

The two mechanisms discussed above are not equally well constrained by our data. The frustum model qualitatively reproduces all three pairwise correlations of Figure 3 when the apical and basal areas are both drawn from the distribution of the experimental area. The quantitative agreement depends on several assumptions that are difficult to assess. The continuity test with the plug flow assumption showed that on the cellular scale and at short timescales there are deviations from volume conservation. The fact that there is volume conservation at long length and time scales may indicate that there is no real mass loss, but the accuracy is not sufficient to rule out real mass loss. The evidence for pulsatile dry-mass excretion is currently indirect: MDCK cells are known to secrete basement-membrane material [51], and only mitotic dry-mass shedding has been measured directly in suspension [33] while the apparent mass loss reported here occurs in cells away from division.

We therefore favour an interpretation in which both effects contribute, with non-prismatic geometry accounting for most of the apparent volume fluctuation and active mass exchange contributing the remainder. A direct test will, for example, require 3D segmentation of refractive index tomograms and simultaneous tracking of apical and basal cell boundaries in addition to methods to analyse extracellular mass build-up.

### C. Cell volume decreases with cell density

Across the confluent density range we sample, the mean cell volume decreases by roughly 25 % while the mean monolayer height increases by a factor of two (Figure 2B,C). These results qualitatively agree with the much larger volume reduction reported by Devany *et al*. [18]. They used the simplifying assumption of a density-independent cell height so that their volume estimates derived from cell area alone systematically misestimate the true volume of cells. Our results agree with their proposed mechanism in which tissue confinement *partially* suppresses cell growth while divisions continue, reducing cell volume successively.

### D. Gamma-shaped height distributions and the pulsation timescale

Once heights are rescaled by their mean, the pixel-wise height distributions collapse onto a gamma-like form across the full range of cell densities studied (Figure 2 and 6). A gamma distribution is the natural outcome of a sum of independent exponentially-distributed increments, and analogous gamma laws are known to describe single-cell protein concentrations [52] and inter-division times [53], both of which are governed by stochastic, burst-like cellular processes. The persistence of a gamma form across cell densities therefore suggests that monolayer height inherits its statistical structure directly from the underlying cell-cycle and growth machinery.

The temporal structure of the height field tells a complementary story. Detrended height autocorrelations (Figure 2H and 6H) have a pronounced negative peak corresponding to oscillations with a ∼ 5 h period at the cellular scale. Two observations make this number meaningful. First, it coincides with the period of velocity and velocity divergence correlations in MDCK monolayers previously reported [9, 37]. This is consistent with the continuity equation that we showed is observed at super-cellular length scale. Second, ∼ 5 h is the same period at which Liu *et al*. [50] observed mass-growth oscillations in single HeLa cells. The coincidence of timescales between intracellular growth oscillations and our tissue-scale height pulsations suggests that the cell-cycle biosynthetic machinery sets the clock that the monolayer pulsations follow, although establishing this causally will require simultaneous measurement of growth rates and height fields.

### E. Propagating modes and the breakdown of plug flow

The structure-factor analysis of the dry-mass field (Appendix A 3, Figure 7) adds a third independent line of evidence that the simple plug-flow picture of an epithelial monolayer is incomplete. The static structure factor exhibits a clear crossover at the cell diameter, *q* ≃ *π/d*_cell_, separating a small-*q* regime in which *S*_*m*_(*q*) ∼ *q*^−2^, characteristic of scale-free density fluctuations and a breakdown of long-range order, from a large-*q* regime *S*_*m*_(*q*) ∼ *q*^−4^ that simply reflects the geometry of individual cells. A Brillouin-like peak whose frequency Ω_*B*_ and half-width Γ_*B*_*/*2 both scale linearly with *q* (Figure 7D), giving an “adiabatic” group velocity of *c*_*s*_ ≃ 15 µm*/*h. Linear *ω*–*q* dispersion is the signature of a propagating elastic mode rather than diffusive relaxation and is naturally accommodated by viscoelastic and active-hydrodynamic descriptions of the monolayer [5, 14–16], but not by the height-averaged plug-flow continuity equation we test in Section III D. The value *c*_*s*_ ≃ 15 µm*/*h is lower than that reported by Zehnder *et al*. [9], but their measurements were taken at lower cell density. We also note the agreement between three different imaging modalities: phase contrast [40], fluorescence intensity proportional to cytosolic mass [9], and direct dry-mass imaging in the present work all yield qualitatively similar structure factors, supporting the interpretations of the monolayer dynamics.

The propagating mode and the breakdown of mass continuity at cellular scales (Section III D) are two facets of the same underlying physics: the monolayer is not a thin, locally incompressible plug-flow fluid but a viscoelastic active medium whose collective excitations carry mass over distances larger than a single cell, and on timescales shorter than mass conservation can be verified locally. Continuum models that aim to describe the full range of observed dynamics must therefore include both elastic restoring stresses and the 3D cell geometry discussed in Section IV C.

### F. What QPI of epithelial monolayers reveals that prior techniques could not

Each of the results in the above is based on a measurement that was not previously accessible for confluent epithelial monolayers. Confocal fluorescence microscopy of membrane labels can resolve cell height with reasonable precision, but it is comparatively slow and phototoxicity inhibits long term, frequent measurements, and does not give direct access to cellular dry mass [8–10, 18]. Stimulated Raman scattering microscopy quantifies protein and lipid concentrations and has been instrumental in establishing the constancy of cellular dry mass concentration of isolated cells and tissues [20, 24], but its acquisition speed and field of view make it ill-suited to resolve the dynamics of collectively migrating and pulsating monolayers. Suspended microchannel resonators provide direct high-precision dry mass of individual cells, but only off the substrate and outside the tissue context [33].

Time-lapse 2D and 3D QPI fills this gap, simultaneously yielding the height field *h*(*x, y, t*), the dry-mass field *m*(*x, y, t*), and segmented cell volumes and masses at minute resolution in the unperturbed monolayer. All three central results, the limited variation in *c*_*d*_ during pulsations (Section III A), the apparent non-conservation of segmented-cell volume at constant *c*_*d*_ (Section III C), and the scale-dependent breakdown of the continuity equation (Section III D), require height and dry mass on the same spatiotemporal grid and could not have been obtained from any single prior technique. The last of these, a direct test of plug-flow continuity against a measured mass field, is to our knowledge new for living epithelia.

These capabilities also place new constraints on previous interpretations. Volume-fluctuation studies based on phase contrast [40], fluorescence intensity [8, 9], or projected cell area [18, 37] have implicitly assumed either that intensity is proportional to cytosolic mass under conserved labelling, or that cell height or volume are constant. Our measurements show that the first assumption is reasonable to first approximation because *c*_*d*_ is regulated on short timescales, but the second is not, because *h*(*ρ*) varies systematically. Finally, our result that cells cannot be assumed to be prisms indicates that interpretations of 2D image segmentations to represent the positions, shapes and velocities of prismatic cells are inaccurate. Combined, these results both support the use of the earlier techniques as proxies for collective motion at the tissue scale and caution against extracting quantitative cell-volume, cell-mass velocity and nematic structure dynamics from them at the cellular scale.

## V. CONCLUSION

Our QPI–based measurements establish that dry mass concentration in MDCK epithelial monolayers is strictly regulated, with a cell density dependent average varying from *c*_*d*_ = 0.1225 at 1300 cells/mm^2^ to *c*_*d*_ = 0.135 at 3300 cells/mm^2^. The dispersion between cells and over time is only 4.5%, despite the large concurrent changes in cell area, height, and projected volume during collective motion. This rules out fluid (water) transport as the dominant driver of height and area pulsations, and challenges interpretations of earlier volume-fluctuation measurements that implicitly attributed volume loss to water efflux. These results rest on simultaneous, time-resolved measurement of cell height, volume, and dry mass in label-free monolayers, a combination that has not been available for previous studies of epithelial collective dynamics.

We find that the mean monolayer height increases approximately linearly with cell density, while mean cell volume decreases, indicating contact inhibition of cell size rather than simple maintenance of constant cell volume. At all densities, monolayer height fields display broad, gamma-like distributions and large spatial and temporal fluctuations, with characteristic oscillation timescales of a few hours that match similar results in the literature. Cell-resolved analysis reveals that height, area, and volume fluctuate synchronously, but with only partial correlations; volume changes are dominated by area fluctuations, and projected volume is not conserved at the single-cell level.

By comparing experiments with simple geometrical models, we demonstrate that frustum cell shapes with constant true volume can reproduce a large part of the observed correlations between apical area, height, and projected volume, implying that 2.5D analyses systematically misestimate cell volume when lateral cell membranes are non-vertical. **The small aspect ratios of the cells suggest that shape changes may not be the complete explanation, a residual signal is consistent with periodic mass exchange between cells and the ECM**. The continuum mass-flux analysis further shows that the height-averaged continuity equation under plug-flow assumptions fails at cellular scales: mass conservation is only recovered after spatial coarse-graining over 2 cell diameters and temporal averaging over 1.6 h. The analysis of the structure factor of mass dynamics shows a propagating elastic mode indicating a viscoelastic active medium whose collective excitations carry mass over distances larger than a single cell.

Taken together, these findings question two widely used assumptions in epithelial monolayer modelling: that cell mass/volume is conserved on the timescales of collective motion, and that monolayers can be treated as 2D sheets with vertical, prism-shaped cells undergoing plug flow. Our results argue that accurate continuum and discrete models of epithelial dynamics must incorporate (i) active regulation of dry mass concentration, (ii) true 3D cell geometry and rearrangements, and (iii) scale-dependent breakdown of volume conservation and plug-flow kinematics. These insights provide a quantitative framework for reinterpreting previous 2D studies and for developing 3D theories and simulations that better capture how mechanical forces, cell packing, and size control jointly shape epithelial behaviour.

## DATA AVAILABILITY

The data that support the findings of this study are available at https://doi.org/10.5281/zenodo.20812823 upon reasonable request. The code used to process and analyse the data is available at https://github.com/siljabl/MDCK-QPI [31].

## ACKNOWLEDGEMENTS

This project has received funding from the European Union’s Horizon 2020 research and innovation programme under the Marie Sklodowska-Curie grant agreement N° 945371. The authors thank UiO:LifeScience for the use of the HoloMonitor and the Hybrid Technology Hub, UiO for the use of the TomoCube, Edna (Xian) Hu for providing the MDCK cells and Vishesh Dubey and Balpreet Singh Ahluwalia for the use of a cell phantom. We thank Sylvain Monnier, Charlotte Riviere, Jean-Paul Rieu, Hèléne Delanoë-Ayari, Claire Desalles, Pierre Recho, Navdeep Rana, Luiza Angheluta-Bauer, and Amin Doostmohammadi for fruitful discussions. We also thank Sumin Lee, Paul Park and Tomocube Inc. for answering our questions about the Tomocube calibration.

## DECLARATION OF INTERESTS

The authors declare no competing interests.

## DECLARATION OF GENERATIVE AI AND AI-ASSISTED TECHNOLOGIES IN THE WRITING PROCESS

During the preparation of this work, the authors used gpt.uio.no and Writefull to improve readability and language. After using this tool, the authors reviewed and edited the content as needed and assume full responsibility for the content of the publication. Appendix B was generated with the help of Claude Opus to summarise the foundational scientific papers underlying the Tomocube analysis. The summary of our understanding and conclusions was communicated to Tomocube Inc. who verified that our understanding was correct. Claude Opus was used to identify inconsistencies that lingered in the manuscript, the corrections of these were performed by the authors.

## CREDIT AUTHOR STATEMENT

**Silja Borring Låstad:** Formal analysis, investigation, software, writing - review and editing. **Nigar Abbasova:** Formal analysis, investigation, software, writing - review and editing. **Thomas Combriat:** Software, writing - review and editing. **Dag Kristian Dysthe:** Conceptualisation, formal analysis, software, writing - original draft, writing - review and editing.

## Appendix A: Supplementary information

### 1. Supplementary movies

**Video S1: Dynamics of refractive index during collective motion at low cell density**. Time-lapse of 3D quantitative phase imaging (QPI) of an MDCK monolayer on glass. The experiment duration is 10 h, acquired at a frame rate of 4 frames/h. The field of view is 567 × 567 µm^2^, and the scale bar corresponds to 100 µm. During the experiment, the cell density is maintained in the range *ρ*_cells_ = 1300–1700 cells/mm^2^.

**Video S2: Dynamics of MDCK height field during collective motion at low cell density**. Height field from same experiment as Video S1.

**Video S3: 3D dynamics of MDCK height field during collective motion at low cell density**. 3D video of height field from same experiment as Video S1, using geometrical proportions that correspond to the actual dimensions of the experiment.

**Video S4: Mass change, mass flux, and their residual in 3D experiment at low density**. Same data and experimental conditions in other movies. The mass change and mass flux are computed within boxes of 10 × 10 µm^2^, then a Gaussian filter with kernel radius 30 × 30 µm^2^ is applied to each frame.

### 2. Disc-wise height data

Some of the height variation in Figure 2D,E comes from intracellular variation, since each cell is the highest at their centre and the lowest at their periphery. To capture the height variation between cells across the monolayer, we performed the same analysis on the Gaussian-filtered height field [22]. The resulting distributions, shown in Figure 6D, are narrower, but still with substantial variation. The relative width at the lowest density is about 30% for the Gaussian-convoluted heights (compared 50% in Figure 2D).

### 3. Structure factor of epithelial mass field

We compute the structure factor *S*_*m*_(**q**, *ω*) = |ℱ(*m*(**r**, *t*)−⟨*m*(**r**, *t*)⟩_**r**_)|^2^, where ℱ(*m*) is the Fourier transform in space and time with wave vector **q** and frequency *ω*. Assuming isotropy, *S*_*m*_(*q, ω*) then represents the azimuthal average of *S*_*m*_(**q**, *ω*). In Figure 7A we show the static structure factor, *S*_*m*_(*q*) = *S*_*m*_(*q*, 0), as function of spatial frequency *q*. There is a clear break in slope at *q* ≃ *π/*10, corresponding to a mean cell diameter of *d*_cell_ = 20 µm. For *q < π/*10 the structure factor scales as *q*^−2^, suggesting scale-free long wavelength fluctuations and breakdown of long range order in the monolayer. At larger wave vectors, the structure factor describes the cell shape and scales as *q*^−4^.

In Figure 7B we plot the dynamic structure factor *S*_*m*_(*q, ω*)*/S*_*m*_(*q*) for several values of *q*. We find two different regimes separated by the crossover *q* = *d*_cell_ described above. For *q < d*_cell_, the dynamic structure factor decays as *ω*^−3^ as found in [40]. For *q > d*_cell_ the decay is markedly slower, and becomes increasingly slow as *q* increases. Following Angelini et al. [40] and Zehnder et al. [9], we model the mass fluctuations as damped harmonic oscillators and fit the sum of Rayleigh and Brillouin shaped peaks to the measured spectra. The fit is illustrated in Figure 7C. In Figure 7D we plot the Brillouin peak frequency Ω_*B*_ and half-width Γ_*B*_*/*2 as function of the spatial frequency *q*. We observe that they both show a propagating mode that scales as *q* with an “adiabatic” group velocity of *c*_*s*_ = Ω_*B*_*/q* = 15 µm/h. This velocity is lower than that reported by Zehnder et al. [9], but their experiments were carried out at lower cell density. We note the agreement of the three different imaging modalities in the analysis above. Angelini et al. [40] analyzed the structure factor of phase-contrast images, Zehnder et al. [9] used fluorescence intensity (which is proportional to cytosolic mass if the fluorescent molecules are conserved), and here we measure dry mass directly by 3D QPI.

### 4. Height, area and volume of frustums and prismatoids

Here, we compute the projected volume of frustum and prismatoid shaped cells, and present the modelling equations used in Figure 4.

If cells are shaped like prisms, their volume can be computed directly from the apical surface area as *V*_prism_ = *V*_projected_ = *h* · *A*_*a*_. In this case, the true cell volume is the same as the projected volume. Like prisms, frustums and prismatoids are geometric shapes that are bounded by two parallel planes. But unlike prisms, their apical area and basal areas do not need to be equal. Prismatoids have an additional degree of freedom compared to that of frustums, in that the number of vertices of the two areas can also differ.

The volume and the height of a frustum are given by

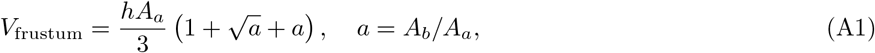

where *A*_*a*_ and *A*_*b*_ are the apical and basal surface areas, respectively. The height of the frustum is then

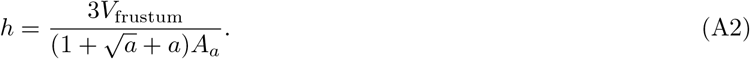

We can then express the projected volume as a function of the frustum volume

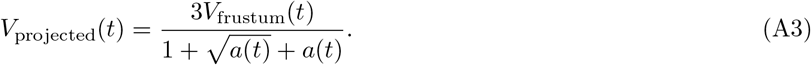

For a constant true volume, we see that the projected volume of a given cell may appear to fluctuate if *a*(*t*) is not constant.

In Figure 4 we have drawn *A*_*a*_ and *A*_*b*_ individually from the experimental distribution of rescaled and detrended cell areas, only accepting combinations that gave *a* ∈ [0.5, 2]. This restriction avoids unrealistic cell shapes. We computed *h* and *V* according to eqs. (A2-A3) and plotted the distributions and correlations of *h, A*_*a*_, and *V*_projected_. The correlations between the variables are 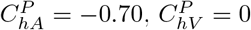, and 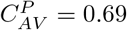.

Likewise, the volume of a prismatoid is

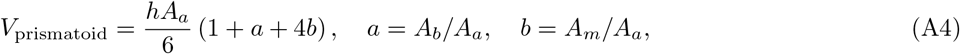

where *A*_*m*_ is the cross-sectional area at the midplane. And we obtain the projected volume and the height as functions of the prismatoid volume

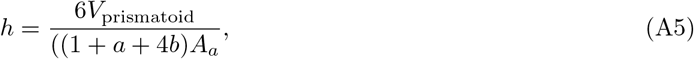

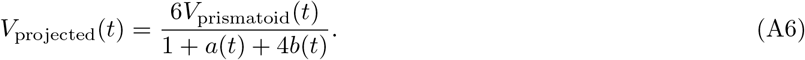

Similar to the case of frustums, we see that the prismatoid cells may appear to have volume fluctuations even if the true volume is constant, if *a*(*t*) or *b*(*t*) is not constant.

The distributions and density plots of prismatoids are shown in Figure 9. We drew *A*_*a*_, *A*_*b*_, and *A*_*m*_ from the experimental distribution of rescaled and detrended cell areas, only accepting combinations that gave *a, b* ∈ [0.5, 2]. The correlations between the variables are 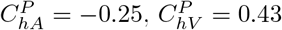, and 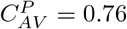.

To test whether the apparent non-conservation of projected cell volume (Section III C) could arise from non-prismatic geometry alone, rather than from genuine changes in cell volume or dry mass, we inverted the frustum model. Assuming a constant true cell volume *V*_0_, the frustum relation (eq. A3) fixes a one-to-one map between a cell’s projected volume and its apical-to-basal area ratio *a* = *A*_*b*_*/A*_*a*_, namely 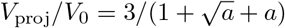. Starting from the measured cell-area distribution (relative standard deviation ≈ 14%), we back-calculated, cell by cell, the distribution of *a* required to reproduce the observed apparent volume fluctuations at constant *V*_0_.

The resulting reconstructed distribution of *a* is right-skewed and approximately Gaussian near its mode, with a relative standard deviation of ≈ 18% and a pronounced tail toward large values, *a* ≈ 1.5–2, i.e. basal areas 1.5–2 times the apical area. Although the central part of this distribution is geometrically admissible, the high-*a* tail is more dubious: for the vertical aspect ratios measured in our monolayers, 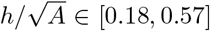, a ratio *a* ≈ 2 would require the lateral cell membranes to be tilted far from vertical (deviating 52^◦^ − 25^◦^ from vertical, for cells with concentric circular areas).

Here, the apical area *A*_*a*_ and basal area *A*_*b*_ are drawn *independently* from the experimental area distribution. This procedure induces a correlation between *A*_*a*_ and the apical–basal area ratio *a*, which in turn generates the correlation between *A*_*a*_ and *V* observed in Fig. 4. In contrast, if we instead draw *a* from the back-calculated distribution described above, and then compute the basal area as *A*_*b*_ = *a* · *A*_*a*_, both the apical area *A*_*a*_ and the ratio *a* become uncorrelated, as do *A*_*a*_ and *V* . In this case, the model fails to reproduce the experimentally measured height and projected-volume distributions. Because it is reasonable to assume that apical and basal areas are at least partially correlated, we infer that deviations from prismatic cell geometry can explain some of the apparent volume fluctuations, but not their full amplitude. The remaining component of the signal likely reflects genuine variation in cell volume and dry mass.

## Appendix B: Tomocube HT-X1 refractive index reconstruction

The Tomocube HT-X1 is a low-coherence holotomography (LC-HT) system that reconstructs a three-dimensional refractive index (RI) distribution from a set of intensity images acquired under varying structured illumination patterns, generated by a digital micromirror device (DMD) placed at the condenser pupil plane. The illumination source is a LED centred at 444 nm. Because the system uses self-interference (no external reference arm), it avoids the coherent noise and vibration sensitivity of conventional Mach–Zehnder interferometry.

### a. Image formation model

Under the *Rytov approximation*—chosen in preference to the Born approximation because the Born approximation systematically underestimates RI and introduces morphological artefacts in biological samples [55]—the normalised log-intensity signal recorded for a given illumination pattern is related linearly to the scattering potential of the sample:

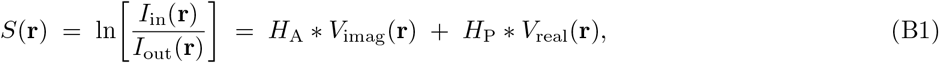

where *I*_in_ and *I*_out_ are the recorded intensities in the *presence* and *absence* of the sample, respectively; *H*_A_ and *H*_P_ are the absorption and phase transfer functions of the microscope (computed analytically from the known pupil intensity distribution); and *V*_real_, *V*_imag_ are the real and imaginary parts of the scattering potential [56]

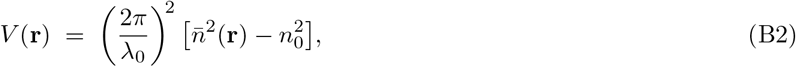

with *λ*_0_ the free-space wavelength and *n*_0_ the RI of the surrounding background medium (a critical input discussed below).

### b. Reconstruction by Wiener deconvolution

The full dataset of *N* illumination patterns yields a system of equations that is solved in the Fourier domain by Wiener (Tikhonov-regularised) deconvolution [55]:

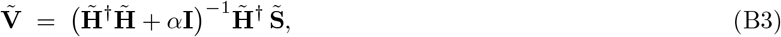

where tildes denote Fourier-transformed quantities and *α* is the regularisation parameter. The real-part RI map is then extracted from *V*_real_ via

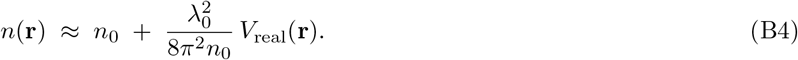

The RI precision of the HT-X1 is quoted as 2 × 10^−4^, characterised by the standard deviation of the reconstructed RI tomogram over a homogeneous sample [57].

### 1. The Missing-Cone Problem and the DC Component

#### a. Origin of the missing cone

In any transmission-geometry microscope, the optical transfer function (OTF) has a characteristic gap in its spatial-frequency support: a double-cone centred on the axial frequency axis (the so-called *missing cone*) is never sampled, regardless of how many illumination angles are used [56]. Physically, this arises because forward-scattered light that propagates nearly parallel to the optical axis carries no additional angle information relative to the unscattered beam. The missing cone predominantly affects low axial spatial frequencies, causing axial elongation and intensity ringing artefacts in the reconstructed RI tomogram; it also reduces the effective axial resolution compared with the lateral resolution [58].

#### b. The DC component and absolute RI

The most severe consequence of the missing cone for absolute RI quantification is its effect on the zero spatial-frequency (DC) component. The DC component of *V*_real_ corresponds to the *spatial average* of the scattering potential across the entire reconstructed volume. Because the DC frequency lies at the very centre of the missing cone, it is **never measured**: the deconvolution in Eq. (B3) has no data with which to constrain the constant offset of *V*_real_, and Wiener regularisation drives this unmeasured component towards zero.

From Eq. (B4), an error Δ*V*_DC_ in the DC component propagates directly as a global, spatially uniform RI offset:

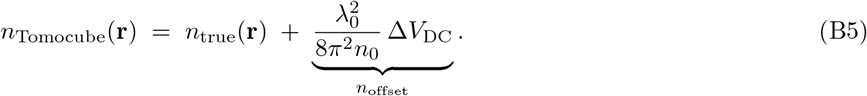

Crucially, *n*_offset_ is *constant across the entire volume*: it shifts every voxel by the same amount, leaving all spatial variations—differences between organelles, between cells, and over time—fully intact. This explains why measured RI contrasts can appear physically reasonable (correct relative ordering, plausible magnitude of spatial heterogeneity) even when the absolute mean RI is clearly incorrect.

#### c. How in-frame background regions normally anchor the DC level

In typical biological experiments, only a fraction of the field of view (FOV) is occupied by cells. The cell-free regions contain only medium, for which *n*(**r**) = *n*_0_ and hence *V*_real_(**r**) = 0 by definition (Eq. (B2)). These background voxels act as an implicit spatial constraint: the reconstruction algorithm, or a post-processing step, can identify that regions with no sample must have *V* = 0 and use this to estimate Δ*V*_DC_ and correct for it. Even without an explicit correction step, the presence of background regions lowers the spatial mean of *V*_real_ in the FOV, effectively pulling the DC estimate closer to zero.

### 2. The Compound Problem for Monolayer Samples

Most published holotomography studies image isolated or sparsely plated cells, organoids embedded in Matrigel domes, or tissue sections—all configurations where a substantial fraction of the FOV contains medium only. Imaging a *confluent cell monolayer* violates this assumption in a structurally important way.

#### a. Absence of in-frame RI reference

When the entire FOV is occupied by cells, two compounding problems arise:

1. **No spatial anchor for the DC level**. The constraint *V* = 0 in background regions cannot be applied because there are no background regions. The only estimate of the DC component comes from the missing-cone extrapolation, which is purely a product of regularisation and therefore unreliable.
2. **Possible error in the background reference image** *I*_out_. From Eq. (B1), the signal *S* requires division by *I*_out_, the intensity recorded without any sample. If *I*_out_ is estimated from a separate calibration frame (dish with medium, no cells), this is in principle correct—but any mismatch between the calibration geometry and the imaging geometry (e.g. different medium depth, slight evaporation, temperature change) introduces a spatially uniform error in *S* and hence a corresponding offset in the reconstructed RI. If, on the other hand, the software infers *I*_out_ from in-frame regions assumed to be cell-free, a confluent monolayer will cause the software to treat cell-covered areas as “background”, forcing *n*_offset_ to absorb the mean RI of the cells rather than that of the medium.

#### b. Impact on RI measurements

The net result is that, for monolayer samples, the absolute mean RI reconstructed by the HT-X1 is undetermined and will reflect the specific circumstances of a given experiment (medium composition, background image quality, FOV coverage by cells) rather than a true physical quantity. Spatial *differences* within and between cells—relative RI between nucleus and cytoplasm, between cell types, and temporal changes within a single timelapse series—remain reliable as long as the missing-cone DC offset is constant within the measurement.

### 3. Relating Tomocube RI to True RI: Linear vs. Offset Model

#### a. The two candidate models

Given the analysis above, we consider two candidate correction models:

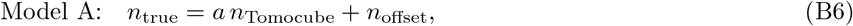

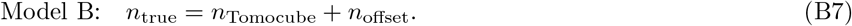

Model B is the special case *a* = 1 of Model A.

#### b. Theoretical basis for Model B (a = 1)

Under the Rytov approximation, the reconstructed RI is (Eq. (B4))

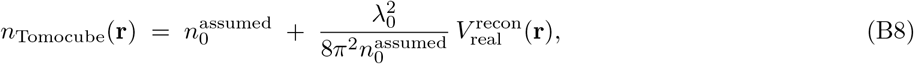

where 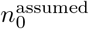 is the value of *n*_0_ used by the software (fixed at instrument configuration), and *V* ^recon^ is the reconstructed scattering potential. For the true RI we have

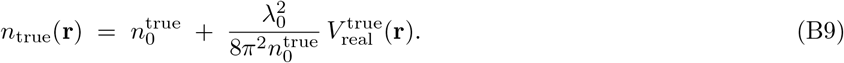

If we assume that the reconstruction faithfully recovers the spatial structure of *V*_real_ (i.e. *V* ^recon^ ≈ *V* ^true^), then the difference between the two expressions is

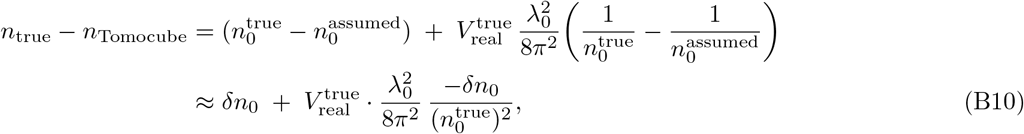

where 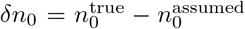. For biological cells, the RI contrast is small: *n*_true_ − *n*_0_ ≲ 0.05, so *V*_real_ ≈ (2*π/λ*_0_)^2^ · 2*n*_0_ · (*n* − *n*_0_) is much smaller than 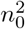, and the second term in Eq. (B10) is of order (*n* − *n*_0_) *δn*_0_*/n*_0_ ≪ *δn*_0_. In the biologically relevant parameter range, the scaling correction is therefore negligible, and the error reduces to a **pure constant offset**:

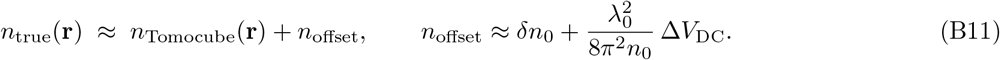

##### Model B is therefore theoretically preferred

A multiplicative factor *a*≠ 1 (Model A) would require either a systematic failure of the Rytov approximation—not expected for the small RI contrasts found in cell biology—or a wavelength-scale calibration error in the OTF (e.g. a wrong NA or wavelength in the software), which would affect all spatial frequencies approximately uniformly and could be checked by inspecting the lateral and axial resolution of known structures.

## Appendix C: Calibration of 2D and 3D QPI

Figure 1 and recent measurements [34] indicate that cell dry mass concentration, *c*_*d*_(*ρ*), increases with cell density, *ρ*. Cell dry mass concentration is proportional to refractive index (RI, *n*) difference: Δ*n*(*ρ*) = *n*(*ρ*) − *n*_0_ = *αc*_*d*_(*ρ*), where *n*_0_ is the medium RI. We use four complementary datasets to fit a model of the refractive index of MDCK monolayers, while calibrating the 2D and 3D QPI measurements:

- 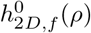: 2D QPI heights measured on fibronectin patches with an absolute zero-height reference in the field of view (FOV). The superscript 0 indicates that these are the initial (uncalibrated) heights, obtained using the initial RI contrast estimate Δ*n*^0^ = 0.04.
- 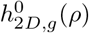: 2D QPI heights measured on glass, without a zero reference in the FOV. These data can be corrected by an additive height offset, Δ*h*, so that they coincide with 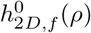 [22]. We refer to the combined dataset as 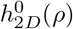.
- *h*_3*D*_(*ρ*): 3D QPI heights obtained from vertical segmentation based on RI contrast [22]. These heights are not affected by the absolute RI offset in the Tomocube measurements.
- Δ*n*_3*D*_(*ρ*): 3D QPI RI contrasts within the monolayer [22]. These contrasts differ from the true values by a dataset-specific additive offset (see Appendix B). In our previous publication [22] we used a per frame offset. After discussion with the manufacturer Tomocube Inc. we have learnt that the offset *n*_offset_ should be kept fixed for a timelapse sequence of frames. Therefore Figure 6 in Låstad et al [22] is not correct.

Our goal is to determine the RI offsets and fit an RI–density model that can be used to correct the 2D QPI heights, *h*_2*D*_(*ρ*), so that they agree with the 3D QPI reference, *h*_3*D*_(*ρ*). For the modeling below, we use the heights averaged over the entire FOV, denoted 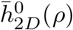 for 2D QPI, and 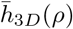 for 3D QPI, and 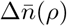 for the RI contrast of 3D QPI.

The calibration is performed in four steps. We first shift 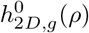 so that it matches 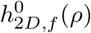. To do this, we fit a second-order polynomial to 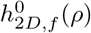 and determine the additive offset that minimizes the deviation between this polynomial and 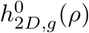. This correction is applied separately for each dish, so that all datasets from a given dish receive the same shift. Next, we shift 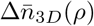 of one 3D dataset so that its trend coincides with that of the other 3D dataset. We then model both the RI contrast and the 3D height as second-order polynomials in density: 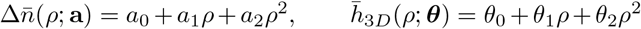. Finally, we correct the constant term 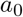 by an additional offset 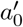 and use the resulting RI model to rescale 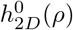 into agreement with *h*_3*D*_(*ρ*). The key quantity is the path integral of the true RI contrast, averaged over the monolayer, 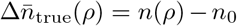, which can be expressed in terms of both 2D and 3D measurements:

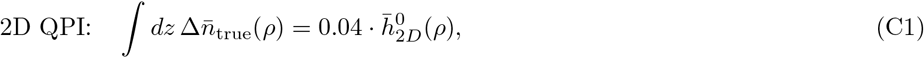

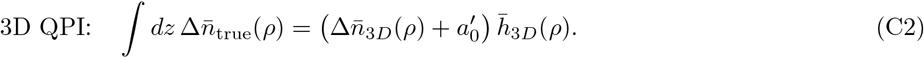

Figure 10B shows *h*_2*D*_(*ρ*) and *h*_3*D*_(*ρ*) after calibration, demonstrating that the 2D QPI heights are brought into agreement with the 3D QPI reference. The resulting calibrated mean RI contrast, 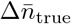, as a function of cell number density is shown in Fig. 10C, and the corresponding dry mass concentration, 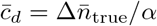, is shown in Fig. 10D.

**FIG. 10.**
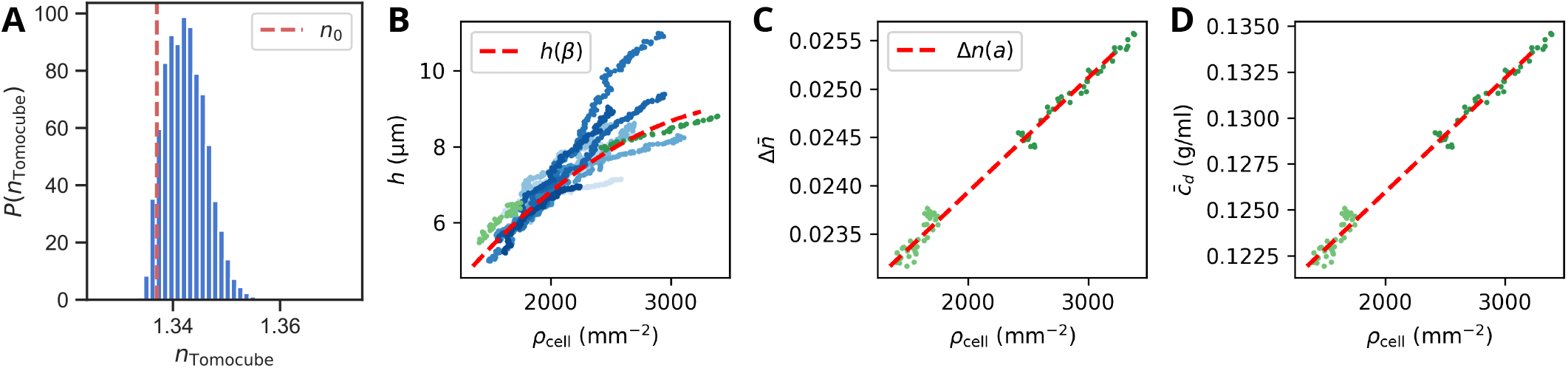
Fitting the refractive index density dependence. **A:** Probability distribution of raw refractive index from one 3D QPI experiment. The reference refractive index of the medium is indicated as *n*_0_. **B:** Mean heights of 2D QPI (blue) and 3D QPI (green) after calibration. The dashed red line shows the fitted model *h*(*ρ*; *β*). **C:** Calibrated mean refractive index 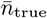 as a function of cell number density after calibration. The dashed red line shows the fitted model Δ*n*(*ρ*; *a*). **D:** Corresponding dry mass concentration 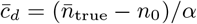 as a function of cell number density.

### 1. Literature summary of MDCK height, volume and dry mass concentration

Both Cadart et al. [17] and Liu et al. [20] report MDCK single cells to have a volume of 2600 µm^3^. Thiagarajan et al. [10] report constant cell volume of 400 µm^3^ (Possibly a typo, a quick back-of-envelope calculation from their figures yield larger volumes) and cell heights within the range 2.5 − 10 µm during pulsations in the monolayer. MDCK height measurements from literature is plotted as function of cell number density in Fig. 11. Volumes are derived from the heights and cell number densities as *V* = *h/ρ*_cell_.

**FIG. 11.**
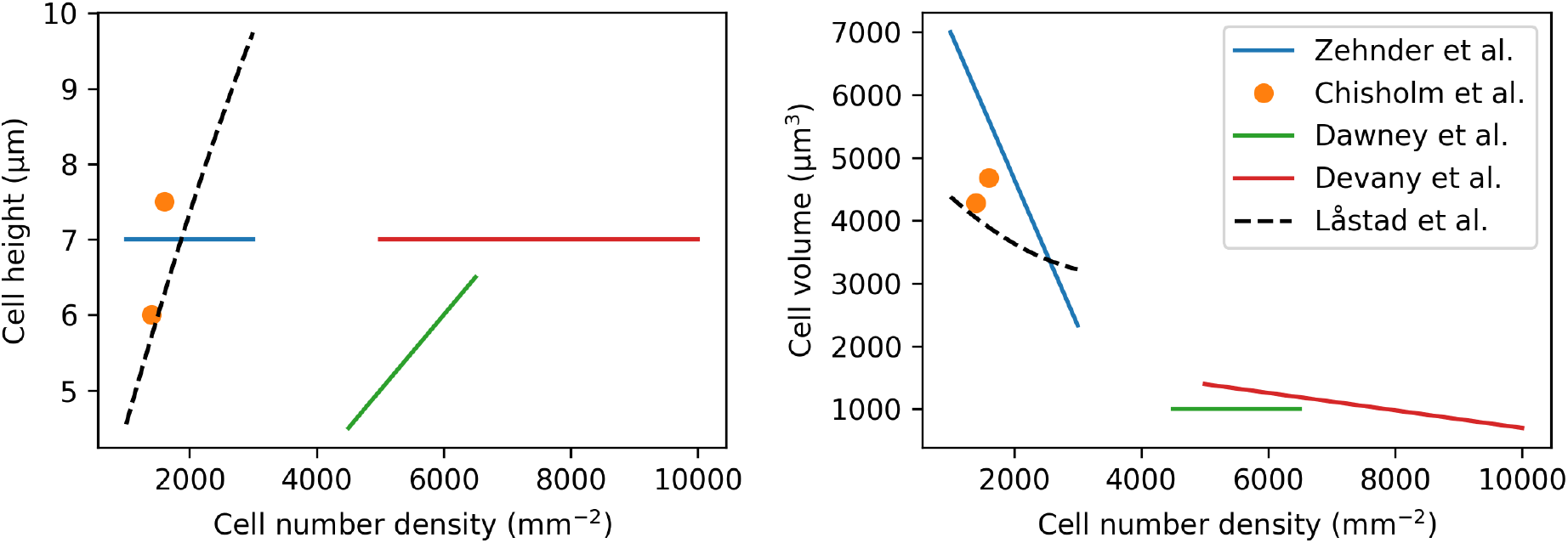
Reported sizes of MDCK. **Left:** Cell heights as function of cell number density. **Right:** Cell volume derived from height and cell number density as *V* = *h/ρ*_cell_.

## Notes

### Competing Interest Statement

The authors have declared no competing interest.

### Summary of Updates

Added section on data availability. Added a few more references.

